# A Domain-general Cognitive Core defined in Multimodally Parcellated Human Cortex

**DOI:** 10.1101/517599

**Authors:** Moataz Assem, Matthew F. Glasser, David C. Van Essen, John Duncan

## Abstract

Numerous brain imaging studies identified a domain-general or “multiple-demand” (MD) activation pattern accompanying many tasks and may play a core role in cognitive control. Though this finding is well established, the limited spatial localization provided by traditional imaging methods precluded a consensus regarding the precise anatomy, functional differentiation and connectivity of the MD system. To address these limitations, we used data from 449 subjects from the Human Connectome Project, with cortex of each individual parcellated using neurobiologically grounded multi-modal MRI features. The conjunction of three cognitive contrasts reveals a core of 10 widely distributed MD parcels per hemisphere that are most strongly activated and functionally interconnected, surrounded by a penumbra of 17 additional areas. Outside cerebral cortex, MD activation is most prominent in the caudate and cerebellum. Comparison with canonical resting state networks shows MD regions concentrated in the fronto-parietal network but also engaging three other networks. MD activations show modest relative task preferences accompanying strong co-recruitment. With distributed anatomical organization, mosaic functional preferences, and strong interconnectivity, we suggest MD regions are well positioned to integrate and assemble the diverse components of cognitive operations. Our precise delineation of MD regions provides a basis for refined analyses of their functions.

## Introduction

Thought and behavior can be conceptualized as complex cognitive structures within which simpler steps are combined to achieve an overall goal (Miller et al. 1960; Luria 1966; Newell 1990). Each step or cognitive episode involves a rich combination of relevant external and internal inputs, computations, and outputs, assembled into the appropriate relations as dictated by current needs. Theoretical proposals have long emphasized that any system capable of such behavior must be equipped with a flexible control structure that can appropriately select, modify and assemble each cognitive step on demand (Norman and Shallice 1986; Duncan et al. 1997; Dehaene et al. 1998; Baddeley 2000; Duncan 2001, 2013; Miller and Cohen 2001; Rigotti 2010).

In line with a system’s role in organizing complex cognition, selective damage to specific regions in the frontal and parietal cortex is associated with disorganized behavior (Milner 1963; Luria 1966; Norman and Shallice 1986), including significant losses in fluid intelligence (Duncan et al. 1995; Glascher et al. 2010; Roca et al. 2010; Woolgar et al. 2010, 2018; Warren et al. 2014). Numerous functional neuroimaging studies converge on a similar set of frontal and parietal regions that are co-activated when performing a diverse range of cognitively demanding tasks, including selective attention, working memory, task switching, response inhibition, conflict monitoring, novel problem solving and many more (Duncan and Owen 2000; Dosenbach et al. 2006; Cole and Schneider 2007; Fedorenko et al. 2013; Hugdahl et al. 2015). We refer to this network as the multiple-demand (MD) system, reflecting their co-recruitment by multiple task demands (Duncan 2010, 2013; Fedorenko et al. 2013). MD activation is commonly reported in lateral and dorsomedial prefrontal cortex, in the anterior insula, and within and surrounding the intraparietal sulcus, with an accompanying activation often reported near the occipito-temporal junction.

Fine-grained activation patterns in MD regions encode many kinds of task-relevant information, including stimulus features, goals, actions, rules and rewards, suggestive of flexible representations shaped by current cognitive requirements (for a recent comprehensive review see (Woolgar et al. 2016)). Consistent with these data from human studies, single-cell studies of putative MD regions in the alert macaque monkey show dynamic, flexible, densely-distributed encoding of information relevant to a current task (Duncan 2001; Miller and Cohen 2001) in which single neurons often show “mixed selectivity”, or nonlinear response to conjunctions of multiple task features (Miller and Cohen 2001; Sigala et al. 2008; Rigotti et al. 2013; Stokes et al. 2013; Fusi et al. 2016; Naya et al. 2017). We and others have proposed that MD regions lie at the heart of cognitive integration, selecting diverse components of cognitive operations across multiple brain systems and binding them together into appropriate roles and relations (Cole and Schneider, 2007; Duncan 2010, 2013; Miller and Cohen 2001; Fusi et al. 2016). Indeed, the MD activation pattern is frequently revealed by studies either employing a task with integrative demands (Prabhakaran et al. 2000) or studies employing a theory-blind search for brain regions with integrative properties, most commonly through indices of connectivity with other brain regions (Power et al. 2013; Shine et al. 2016; Gordon et al. 2018).

While MD activation has been reported since the early days of human brain imaging (Duncan and Owen 2000), a consensus is lacking over five core questions. (i) What is the precise extent and topography of MD regions in human cortex and their relation to other immediately adjacent regions that have very different functional properties (e.g. see Fedorenko et al. 2012)? (ii) What is the degree of functional differentiation within the MD network? There are many rival proposals and little agreement across studies (Champod and Petrides, 2010; Dosenbach et al., 2007; Hampshire et al., 2012; Lorenz et al., 2018; Yeo et al., 2015). (iii) What is the precise relationship to “canonical” resting-state fMRI (rfMRI) brain networks revealed by various ways of grouping regions based on the strength of their time-series correlations? A “fronto-parietal network” (FPN) shows strong anatomical similarity with MD activations (Power et al. 2011; Yeo et al. 2011; Blank et al. 2014; Laumann et al. 2015; Ji et al. 2019), but a finer examination of its overlap with MD activations and relations with other networks is currently lacking. (iv) What are the links – long suspected but rarely examined in detail – with accompanying MD activation in regions of the basal ganglia, thalamus and cerebellum (Buckner et al. 2011; Choi et al. 2016; Halassa and Kastner 2017)? (v) What are the correspondences with putative cortical MD regions identified in other primates (Ford et al. 2009; Mitchell et al. 2016; Premereur et al. 2018)?

Our understanding of these and other aspects of MD function will surely benefit from improved anatomical localization. MD activation has often been described in terms of large, loosely-defined regions such as “dorsolateral prefrontal cortex” that also include regions having very different functional responses and sharp transition boundaries (Glasser, Coalson, et al. 2016). Traditional fMRI analysis methods typically use non-optimal inter-subject registration and apply substantial smoothing, both of which blur across functional boundaries. While problems of this sort may be offset by individual-subject region of interest (ROI) methods, for many questions consensus ROIs are lacking, limiting comparison and integration of results across studies.

To address these issues, we turned to the large-scale data and novel analysis approach of the Human Connectome Project (HCP). To improve delineation of functional regions, HCP analyses used high quality multimodal MRI features (cortical thickness, myelin content, rfMRI connectivity, task fMRI activation), along with surface-based analysis methods (Glasser et al. 2013; Glasser, Smith, et al. 2016; Coalson et al. 2018) and new areal-feature-based registration algorithms (Robinson et al. 2014, 2018). Here we relate MD activation to the state-of-the-art multi-modal HCP parcellation of human cortex into 360 regions (180 per hemisphere), in which areal delineations were derived using overlapping multi-modal criteria, and areas were named to reflect correspondences with the neuroanatomical literature.

We analyzed data from 449 HCP subjects, each having a defined individual-specific cortical parcellation. Our analysis was based on three suitable fMRI task contrasts available in the HCP data: working memory 2-back versus 0-back (WM 2bk>0bk), hard versus easy relational reasoning (Relational H>E), and math versus story (Math>Story). The first two are standard hard>easy contrasts as commonly used to define MD activation (Duncan and Owen, 2000; Fedorenko et al., 2013; e.g. for n-back MD activation: Gray et al., 2003; Owen et al., 2005; e.g. for reasoning MD activation: Duncan, 2000; Watson and Chatterjee, 2012). Math>Story was added because previous results show a strong MD-like activation pattern associated with arithmetic processing (Amalric and Dehaene 2016, 2017). For working memory and relational reasoning, stimuli were visual, whereas for Math>Story, stimuli were auditory. The other four HCP tasks lacked typical MD contrasts and were not used. Combining data from the 3 task contrasts, we determined which areas show MD properties and examined their functional profiles, patterns of resting state connectivity, and relations to subcortical structures.

Our results reveal an extended, largely symmetrical MD network of 27 cortical areas, distributed across frontal, parietal and temporal lobes. We divide this extended MD system into a core of 10 regions most strongly activated and strongly interconnected, plus a surrounding penumbra, and we relate this functional division to canonical resting state networks also derived from HCP data (Ji et al. 2019). Across the extended MD system, activation profiles for our 3 task contrasts suggest a picture of substantial commonality, modulated by modest but highly significant functional differentiations. MD activation, and strong functional connectivity with the cortical MD core, are also identified in several subcortical regions. Our results define a highly specific, widely distributed and functionally interconnected MD system, which we propose forms an integrating core for complex thought and behavior.

## Materials and Methods

### 1. Subjects

The analyzed dataset consisted of 449 healthy volunteers from the Human Connectome Project (HCP) S500 release. Subjects were recruited from the Missouri Twin Registry (186 males, 263 females), with age ranges 22-25 (n=69), 26-30 (n=208), 31-35 (n= 169), and 36+ (n=3). Informed consent was obtained from each subject as approved by the institutional Review Board at Washington University at St. Louis.

### 2. Image Acquisition

MRI acquisition protocols have been previously described (Glasser et al. 2013; Smith et al. 2013; Uǧurbil et al. 2013). All 449 subjects underwent the following scans: structural (at least one T1w MPRAGE and one 3D T2w SPACE scan at 0.7 mm isotropic resolution), rfMRI (4 runs X 15 minutes), and task fMRI (7 tasks, 46.6 minutes total). Images were acquired using a customized 3T Siemens ‘Connectom’ scanner having a 100mT/m SC72 gradient insert and using a standard Siemens 32-channel RF receive head coil. Whole brain rfMRI and task fMRI data were acquired using identical multi-band EPI sequence parameters of 2 mm isotropic resolution with a TR=720 ms. Spin echo phase reversed images were acquired during the fMRI scanning sessions to enable accurate cross-modal registrations of the T2w and fMRI images to the T1w image in each subject (standard dual gradient echo fieldmaps were acquired to correct T1w and T2w images for readout distortion). Additionally, the spin echo field maps acquired during the fMRI session (with matched geometry and echo spacing to the gradient echo fMRI data) were used to compute a more accurate fMRI bias field correction and to segment regions of gradient echo signal loss.

### 3. Task Paradigms

Each subject performed 7 tasks in the scanner over two sessions. In the current study we analyzed data from 3 tasks: working memory (performed in session 1), math/language and relational reasoning (performed in session 2). Subjects performed 2 runs of each task. The following task details are adapted from Barch et al. (2013) on HCP fMRI tasks.

#### Working Memory

Each run consisted of 8 task blocks (10 trials of 2.5 s each, for 25 s) and 4 fixation blocks (15 s each). Within each run, 4 blocks used a 2-back working memory task (respond ‘target’ whenever the current stimulus was the same as the one two back) and the other 4 used a 0-back working memory task (a target cue was presented at the start of each block, and a ‘target’ response was required to any presentation of that stimulus during the block). A 2.5 s cue indicated the task type (and target for 0-back) at the start of the block. On each trial, the stimulus was presented for 2 s, followed by a 500 ms ITI. In each block there were 2 targets, and (in the case of the 2-back task) 2–3 non-target lures (repeated items in the wrong n-back position, either 1-back or 3-back). Stimuli consisted of pictures of faces, places, tools and body parts; within each run, the 4 different stimulus types were presented in separate blocks. Subjects had to respond to non-targets using a middle finger press and to targets using an index finger press.

#### Math/language

Each run consisted of 4 blocks of a math task interleaved with 4 blocks of a story task. The lengths of the blocks varied (average of approximately 30 s), but the task was designed so that the math task blocks matched the length of the story task blocks, with some additional math trials at the end of the task to complete the 3.8 min run as needed. The math task required subjects to complete addition and subtraction problems, auditorily presented. Each trial had a problem of the form “X + Y =” or “X – Y =”, followed by two choices. The subjects pushed a button to select either the first or the second answer. Problems were adapted to maintain a similar level of difficulty across subjects. The story blocks presented subjects with brief auditory stories (5–9 sentences) adapted from Aesop’s fables, followed by a 2-alternative forced choice question that asked the subjects about the topic of the story. The example provided in the original Binder paper (p. 1466) is “For example, after a story about an eagle that saves a man who had done him a favor, subjects were asked, ‘That was about revenge or reciprocity?’”. For more details on the task, see (Binder et al. 2011).

#### Relational Reasoning

Stimuli were drawn from a set of 6 different shapes filled with 1 of 6 different textures. In the hard condition, subjects were presented with 2 pairs of objects, with one pair at the top of the screen and the other pair at the bottom of the screen. They were told that they should first decide what dimension(s) differed across the top pair of objects (shape or texture) and then they should decide whether the bottom pair of objects also differed along the same dimension(s) (e.g., if the top pair differs only in shape, does the bottom pair also differ only in shape?). In the easy condition, subjects were shown two objects at the top of the screen and one object at the bottom of the screen, and a word in the middle of the screen (either “shape” or “texture”). They were told to decide whether the bottom object matched either of the top two objects on that dimension (e.g., if the word is “shape”, is the bottom object the same shape as either of the top two objects?). Subjects responded with their right hand, pressing one of two buttons on a handheld button box, to indicate their response (“yes” or “no”). For the hard condition, stimuli were presented for 3500 ms, with a 500 ms ITI, with four trials per block. In the easy condition, stimuli were presented for 2800 ms, with a 400 ms ITI, with 5 trials per block. Each type of block (hard or easy) lasted a total of 18 s. In each of the two runs of this task, there were 3 hard blocks, 3 easy blocks and 3 16 s fixation blocks.

### 4. Data preprocessing

Data were preprocessed using the HCP’s minimal preprocessing pipelines (Glasser et al. 2013). Briefly, for each subject, structural images (T1w and T2w) were corrected for spatial distortions. FreeSurfer v5.3 was used for accurate extraction of cortical surfaces and segmentation of subcortical structures. To align subcortical structures across subjects, structural images were registered using non-linear volume registration to Montreal Neurological Institute (MNI) space.

Functional images (rest and task) were corrected for spatial distortions, motion corrected, and mapped from volume to surface space using ribbon-constrained volume to surface mapping. Subcortical data were also projected to the set of extracted subcortical structure voxels and combined with the surface data to form the standard CIFTI grayordinates space. Data were smoothed by a 2mm FWHM kernel in the grayordinate space that avoids mixing data across gyral banks for surface data and avoids mixing areal borders for subcortical data. Rest and task fMRI data were additionally identically cleaned up for spatially specific noise using spatial ICA+FIX (Salimi-Khorshidi et al. 2014) and global structured noise using temporal ICA (Glasser et al. 2018).

For accurate cross-subject registration of cortical surfaces, a multi-modal surface matching (MSM) algorithm (Robinson et al. 2014) was used to optimize the alignment of cortical areas based on features from different modalities. MSMSulc (‘sulc’: cortical folds average convexity) was used to initialize MSMAll, which then utilized myelin, resting state network (RSN) and rfMRI visuotopic maps. Myelin maps were computed using the ratio of T1w/T2w images (Glasser and Van Essen 2011; Glasser et al. 2014). Individual subject RSN maps were calculated using a weighted regression method (Glasser, Coalson, et al. 2016).

### 5. HCP multi-modal parcellation and areal classifier

The HCP multi-modal parcellation map (MMP) 1.0 (Glasser, Coalson, et al. 2016) was first created using a semi-automated approach utilizing the group average maps of multiple modalities (cortical thickness, myelin, resting state functional connectivity, and task activations). For each modality, the gradient was computed as the 1^st^ spatial derivative along the cortical surface; ridges were local regions with the highest value and thus the most sudden change in a feature. Overlapping gradient ridges across modalities were used to draw putative areal borders with manual initialization and algorithmic refinement. Defined areas were reviewed by neuroanatomists, compared whenever possible to previously identified areas in the literature, and labelled. This resulted in defining 180 areas per hemisphere. A multi-modal areal classifier was then used for automated definition of areas in each subject using the multi-modal feature maps. The classifier was trained, tested and validated on independent groups of subjects from the same 449 cohort used in this study (Glasser, Coalson, et al. 2016).

### 6. Task fMRI analysis

Task fMRI analysis steps are detailed in Barch et al. (2013). Briefly, autocorrelation was estimated using FSL’s FILM on the surface. Activation estimates were computed for the preprocessed functional time series from each run using a general linear model (GLM) implemented in FSL’s FILM (Woolrich et al. 2001). For the *working memory* task, 8 regressors were used - one for each type of stimulus in each of the N-back conditions.

Each predictor covered the period from the onset of the cue to the offset of the final trial (27.5 s). For the *math* task, 2 regressors were used. The math regressor covered the duration of a set of math questions designed to roughly match the duration of the story blocks. The story regressor covered the variable duration of a short story, question, and response period (∼30 s). For the *relational reasoning* task, two regressors were used, each covering the duration of 18 s composed of four trials for the hard condition and five trials for the easy condition. In each case, linear contrasts of these predictors were computed to estimate effects of interest: WM 2bk>0bk, Relational H>E, and Math>Story.

All regressors were convolved with a canonical hemodynamic response function and its temporal derivative. The time series and the GLM design were temporally filtered with a Gaussian-weighted linear highpass filter with a cutoff of 200 seconds. Finally, the time series was prewhitened within FILM to correct for autocorrelations in the fMRI data. Surface-based autocorrelation estimate smoothing was incorporated into FSL’s FILM at a sigma of 5mm. Fixed-effects analyses were conducted using FSL’s FEAT to estimate the average effects across runs within each subject.

For further analysis of effect sizes, beta ‘cope’ maps were generated using custom built MATLAB scripts after moving the data from the CIFTI file format to the MATLAB workspace and after correction of the intensity bias field with an improved method (Glasser et al 2016a). Activation estimates on cortical surface vertices were averaged across vertices that shared the same areal label in a given subject. Unless mentioned otherwise, parametric statistical tests (one-sample and paired sample t-tests) were used.

### 7. rfMRI Functional connectivity analysis

For each subject, a ‘parcellated’ functional connectivity (FC) map was computed by averaging the time series across cortical vertices that shared the same areal label and correlating the average time series, giving a 360×360 cortical FC matrix for each subject.

For comparison of connection types (Figure 3b, Figure 4b), connectivities for each subject were simply averaged across each group of areas following r-to-z transformation. 1-r was used as the distance measure for the multi-dimensional scaling analysis (MATLAB function cmdscale).

Subcortical analysis was based on the group average dense FC maps for a split-half division of the subjects (210P and 210V; the parcellation and validation groups used in Glasser, Coalson, et al. 2016). For each subcortical voxel, an average connectivity to the cortical MD core was obtained by first calculating FC with each core area (after averaging across each area’s vertices), and then averaging these connectivities following r-to-z transformation. A permutation testing approach (100,000 permutations) was used to identify significant voxels by building a null distribution for each voxel based on its FC estimate to sets of 10 randomly selected cortical areas across both hemispheres. A voxel was determined as significantly connected to the MD system when its FC estimate was in the top 97.5^th^ percentile.

### Data availability

Data used for generating each of the imaging-based figures are available on the BALSA database (https://balsa.wustl.edu/study/B4nkg). Selecting the URL at the end of each figure will link to a BALSA page that allows downloading of a scene file plus associated data files; opening the scene file in Connectome Workbench will recapitulate the exact configuration of data and annotations as displayed in the figure.

## Results

### Cortical organization of the MD system at the group level

We analyzed a cohort of 449 HCP subjects **(for details on data acquisition and preprocessing see Methods sections 1-4).** For an initial overview of the MD activation pattern, we calculated a group average MD map by averaging the group average beta maps of the 3 task contrasts and overlaying the resulting combined map on the HCP MMP 1.0 parcellation areal borders **(see Figure S1 for each contrast separately).** Group average maps were generated by aligning each subject’s multi-modal maps using areal-feature-based surface registration (MSMAll, Robinson et al 2014; 2018**; see Methods section 4).** MSMAll registration is initialized by cortical folding patterns and then uses myelin and connectivity features to significantly improve the alignment of areas across subjects (Coalson et al. 2018), thus allowing us to identify cortical areas most strongly overlapping with MD activations.

The resulting overview is shown on left and right inflated cortical surfaces in **Figure 1a**, and on a cortical flat map of the left hemisphere in **Figure1b**. The results highlight 9 patches of activation distributed across the cortical sheet. On the lateral frontal surface are four clearly distinct patches that show strong bilateral symmetry, with surrounding inactive regions: a dorsal region (patch 1), a premotor region (patch 2), a mid-frontal region (patch 3) and a frontal pole region (patch 4). Patch 5 is delineated in and surrounding the anterior insula. Tight bands of MD activation are also identifiable in dorsomedial frontal cortex (patch 6), along the depths of the intraparietal sulcus spreading up to the gyral surface (patch 7), and in dorsomedial parietal cortex (patch 8).

**Figure 1.**
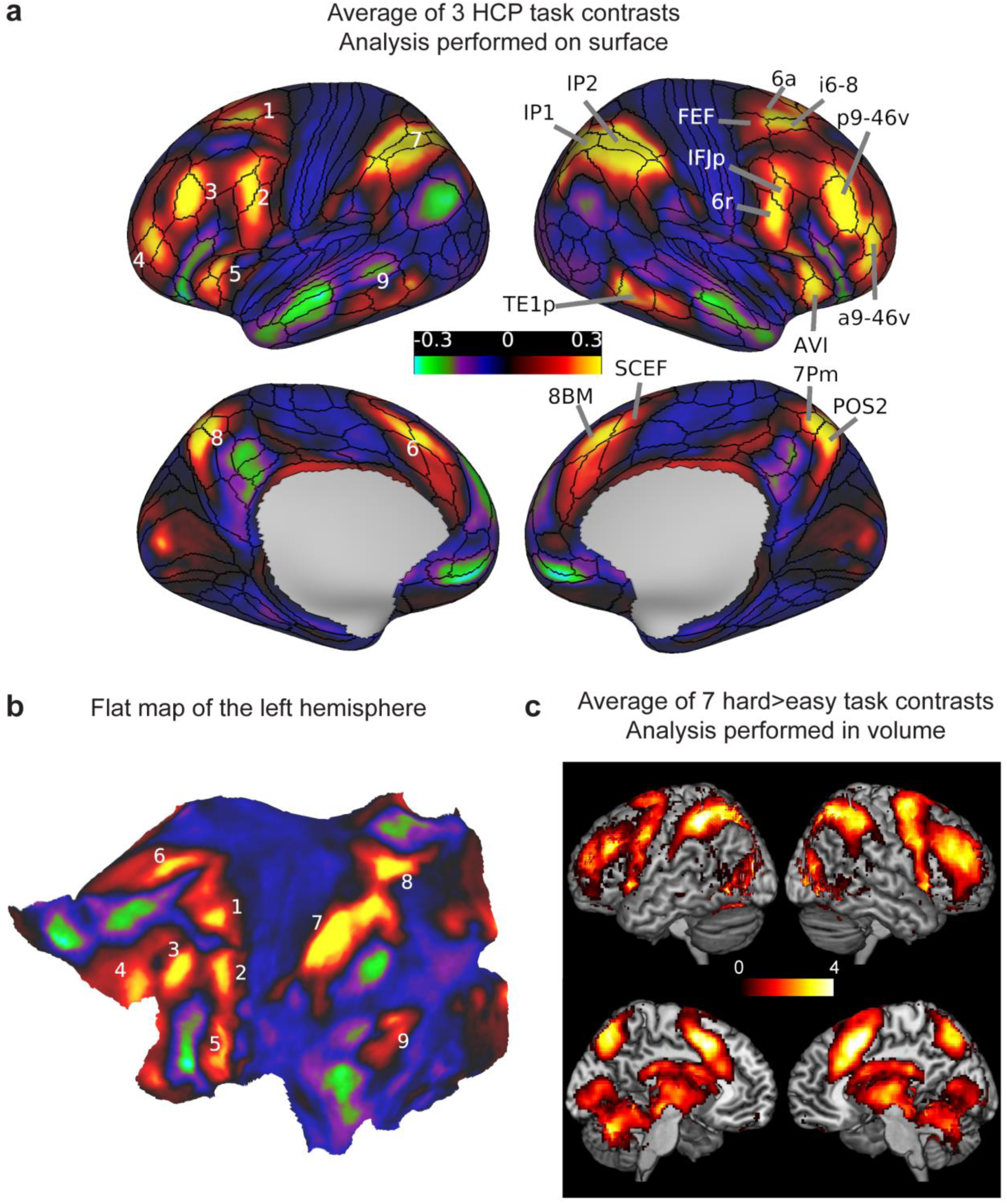
**(a)** Average of the 3 HCP group average task contrasts (WM 2bk>0bk, Relational H>E, Math>Story). Values are beta estimates. Black contours correspond to the HCP multi-modal parcellation MMP_1.0 (210V) areal borders. Numbers on the left hemisphere correspond to visually separable patches of activity distributed across the cortex. **(b)** The same activity of the left hemisphere projected on a flattened cortical sheet. Numbers correspond to the same patches labelled in (a). **(c)** Volumetric MD map from Fedorenko et al. (2013) computed by averaging 7 hard>easy task contrasts (2mm smoothed) displayed on a volume rendering of lateral surface (above) and medial slice (below) of the MNI template. Values are t-statistics. Data available at http://balsa.wustl.edu/lL9nj

The MD region often reported near the occipito-temporal border is also evident in posterior temporal cortex (patch 9). The right hemisphere view in **Figure 1a** identifies cortical areas showing the strongest MD activations.

For comparison, **Figure 1c** shows a previous MD group-average volumetric map generated from the conjunction of 7 hard>easy task contrasts (Fedorenko et al. 2013). Though the two maps are broadly similar, this comparison highlights the improved definition obtained with the HCP data and surface-based and areal-feature-based registration methods. Even based on these average data, the improved co-registration of the HCP data allows clearer delineation of functional regions, as predicted by Coalson et al., 2018. Rather than broad, fuzzy swaths of MD activation, these data provide evidence for a more tightly localized, though anatomically distributed network of MD regions.

### Definition of extended and core MD regions using subject-specific cortical parcellation

For our primary analysis, each subject’s cerebral cortex was parcellated into 360 regions (180 per hemisphere) corresponding to the HCP Multi-Modal Parcellation (MMP) 1.0. Parcellation used an automated classifier to define the borders of each area based on learned features from multiple MRI modalities, including cortical thickness, myelin content, rfMRI connectivity and task fMRI activations **(see Methods section 5)**. Subject-specific parcellation ensured that task and rest fMRI signals extracted from the defined areas would respect individual differences in their sizes, shapes and locations even in the case of subjects having atypical topologic arrangements. We averaged beta values across vertices within each area, yielding one value per area per subject. For each of our 3 behavioral contrasts, we identified areas with a significant positive difference across the group of 449 subjects (p<0.05, Bonferroni corrected for 180 areas). Given largely bilateral activation **(Figure 1),** to improve signal-to-noise ratio (SNR) and statistical power we averaged areal activations across hemispheres.

The conjunction of significant areas across the 3 contrasts revealed a set of twenty-seven areas, which we refer to as the extended MD system **(Figure 2a**; note that average activations from the two hemispheres are projected onto the left). The distribution of the areas closely matches the activations observed in **Figure 1a** and has broad similarity to previous characterizations of MD activation but with substantially improved anatomical precision and several novel findings.

**Figure 2.**
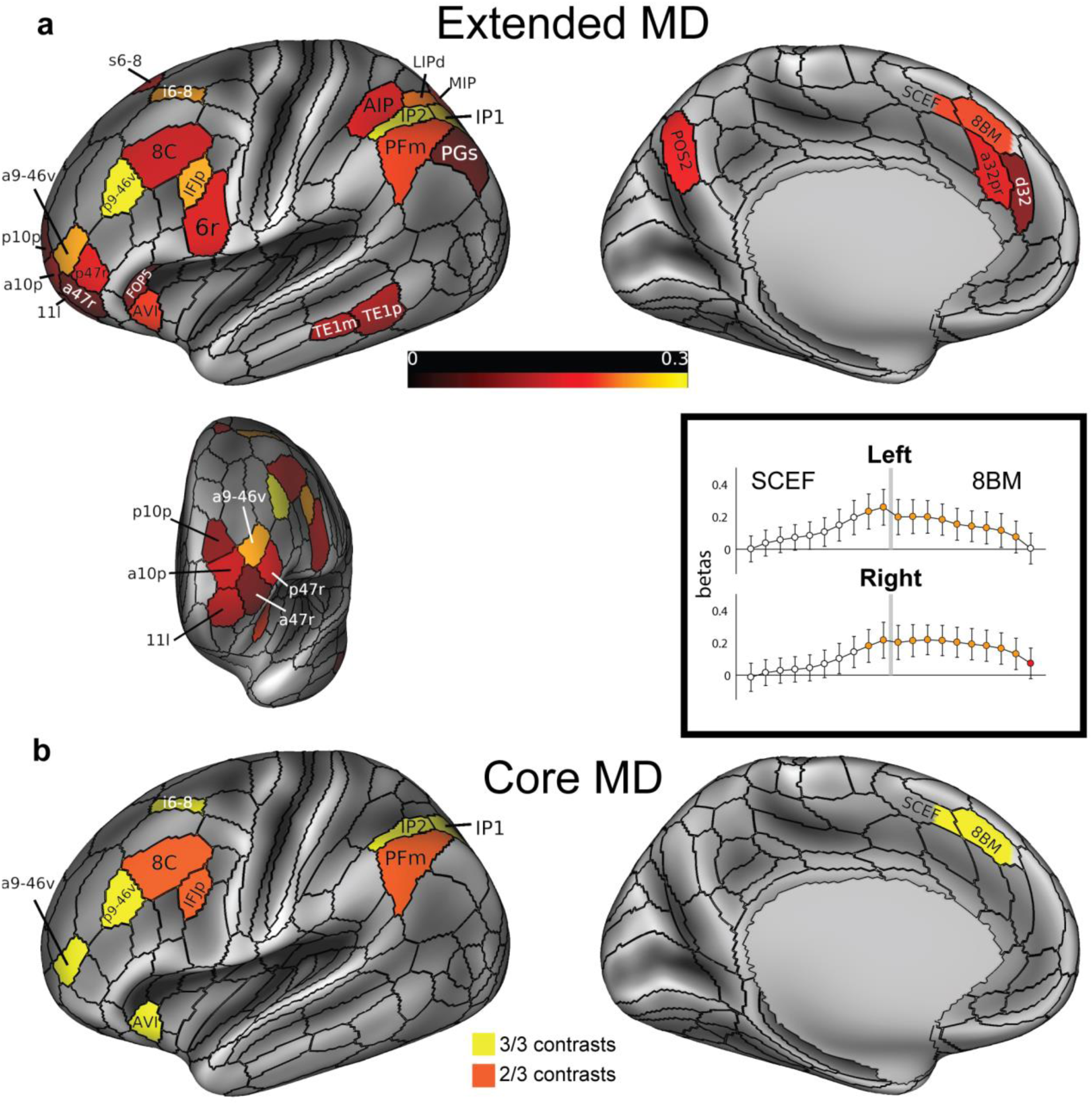
**(a)** The extended MD system: conjunction of significant areas across 3 functional contrasts. Areal colors reflect average beta values across the 3 contrasts analyzed in relation to subject-specific parcellations. Data are averaged across hemispheres, and for illustration projected here onto the left lateral and medial surfaces (*top*) and an anterior view of frontal pole parcels (*bottom left*). Box (*bottom* right) displays pattern of activity in regions SCEF (posterior) and 8BM (anterior), divided into posterior to anterior segments in relation to subject-specific parcellations. Grey bar indicates 8BM/SCEF border. Orange indicates segments that are part of the extended MD system when activity from both hemispheres is combined (i.e. segments with activity significantly above zero in all 3 behavioral contrasts). Red indicates one additional segment that survives as part of the extended MD system when activity from each hemisphere is tested separately. **(b)** The core MD system: areas with activity estimates that were significantly higher than the mean activity of all extended MD areas in all 3 contrasts (yellow) and 2 out of 3 contrasts (orange). Data available at http://balsa.wustl.edu/qNLq8

On the dorsal lateral frontal surface, we identify area i6-8 which is immediately anterior to area FEF (a common assignment for activations in this region). i6-8 is a newly defined area in the HCP MMP1.0, in the transitional region between classical BA6 and BA8. Localization of MD activation in i6-8, rather than FEF, suggests distinctness from activations driven simply by eye movements in complex tasks. In the HCP MMP1.0, FEF is clearly defined as a distinct area from i6-8 based on several criteria including its location as a moderately myelinated patch just anterior to the eye-related portion of the motor cortex and its strong functional connectivity with the LIP/VIP visual complex and the premotor eye field area (PEF) (Glasser et al., 2016).

Near the frontal pole, we identify area a9-46v as a strongly active MD region, separated from the posterior region p9-46v. This separation confirms prior indications of a distinct anterior MD frontal region **(see Figure 1c).** Both a9-46v and p9-46v areas overlap with area 9-46v as delineated cyto-architectonically by Petrides and Pandya (1999) but here are separated into anterior and posterior portions by intervening areas 46 and IFSa that differ in their myelin and functional connectivity profiles (Glasser et al., 2016). Posterior to p9-46v is a further focus of activation in IFJp, with weaker activation in the surrounding regions 8C and 6r.

In the anterior insula, we identify AVI and an adjacent region of the frontal operculum, FOP5. AVI overlaps with superior portions of the architectonic area Iai of Öngür et al., 2003 (see Glasser et al., 2016). Previous work has attempted to distinguish activation in the anterior insula from the adjacent frontal operculum, with the peak often near the junction of the two (Amiez et al. 2016). In our data, AVI is the more strongly activated.

While previous characterizations of parietal MD activation have focused on the intraparietal sulcus broadly, our results reveal a more detailed picture, with strongest MD activation in intraparietal sulcus areas IP1 and IP2, bordered by relatively weaker MD areas dorsally (AIP, LIPd, MIP) and ventrally (PFm, PGs). In dorso-medial parietal cortex, there have been previous indications of an additional MD region (see **Figure 1c**). Here we robustly assign this mainly to area POS2, a newly defined MMP1.0 area that differs from its neighbors in all major multi-modal criteria.

On the lateral surface of the temporal lobe we identify two further MD areas, TE1m and TE1p. In many previous studies, fronto-parietal MD activation has been accompanied by a roughly similar region of activity in temporo-occipital cortex (e.g. Fedorenko et al., 2013). In many cases, a reasonable interpretation would be higher visual activation, reflecting the visual materials of most imaging studies. In the current study, however, the arithmetic task was acoustically presented, whereas the other two contrasts were visual, suggesting a genuine MD region.

In **Figure 1a**, the dorso-medial frontal activation spans the border between 8BM/SCEF. In the individual-subject analysis, however, SCEF was not significantly activated across all 3 contrasts. We thus investigated whether the activation indeed spans the border between the two areas. For each subject-specific areal definition, we divided each of the two areas into 10 equal segments along their anterior to posterior extent. **Figure 2a** shows that activation in this region starts to build up midway along SCEF, peaks at the border and is sustained throughout 8BM. We then tested whether each segment would survive as an extended MD region on its own. Indeed, all 8BM segments (except for the one most anterior segment on the left hemisphere) survived, whereas only the anterior 2 segments of SCEF were statistically significant (**Figure 2a**; see **Figure S2** for further independent evidence of heterogeneity around the 8BM/SCEF border). Based on these results, for subsequent analyses we combined the statistically significant segments of 8BM and SCEF into a single ‘area’ labelled 8BM/SCEF.

To evaluate the reliability of our results, we identified extended MD regions after splitting our subjects into two independent groups constructed to avoid shared family membership (210P and 210V, the parcellation and validation groups, respectively, used to create the HCP MMP1.0 in Glasser et al., 2016). Using similar criteria as for **Figure 2a** (i.e., conjunction of 3 positive contrasts across the group of 210 subjects, each contrast p<0.05 Bonferroni corrected for 180 areas), we identified 24 out of 27 regions in the 210P group (missing regions: 6r, AIP, FOP5) and 25 regions in the 210V group (missing regions: a47r, AIP). No additional regions were identified in either group. Thus for the remainder of the analysis, we retained the full set of 27 regions based on the complete data set.

To delineate more precisely the most active areas within the extended MD system, for each contrast we identified areas with activation stronger than the mean across the full set of 27 regions (one sample t-test, p<0.05, Bonferroni correction for the 27 extended MD areas). Seven areas were significant in all three contrasts: i6-8, p9-46v, a9-46v, combined 8BM/SCEF area, AVI, IP2 and IP1. Three more areas were significant in two of the three contrasts (**Figure 2b**): IFJp (relational reasoning and math), 8C and PFm (working memory and relational reasoning). We refer to this group of 10 areas as the core MD system, with remaining areas of the extended MD system termed the MD penumbra.

Though our main analysis used individual-specific cortical parcellations, we wondered how well results would replicate using just the group-average parcellation. For most areas, previous work shows that the areal-fraction of individually defined parcels captured by group-defined borders reaches 60%-70% (Coalson et al. 2018). To investigate this question, we repeated our analysis using the HCP_MMP1.0 group average parcellation. As expected, using group-defined regions, we identified the same set of 27 MD regions, plus 4 more (areas 44, IFJa, 9-46d, 7Pm). While individual-specific parcellations likely provide the best available areal delineation, for many purposes the group-defined cortical parcellation may be sufficient.

Overall, these results identify an extended set of domain-general MD regions. Using HCP data and analysis allowed the identification of several novel MD areas and improved localization of previously reported ones. In the following sections, we further explore the functional properties of the 27 core and penumbra regions.

### Functional connectivity of the multiple-demand cortex and its relation to resting-state networks

To investigate functional connectivity (FC) patterns within the MD network and in relation to the rest of the brain, a FC matrix for each subject was calculated (180×180 areas per hemisphere; full correlation of spatial ICA+FIX and temporal ICA-cleaned time series; **see Methods section 7**). In this analysis, we retained the original 8BM and SCEF parcellation, considering 8BM as core and SCEF as penumbra.

**Figure 3a** shows the group average connectivity matrix for the extended MD system, separated into core and penumbra. Despite their wide spatial separation, core MD areas show stronger functional connectivity with each other than with the penumbra. To test the robustness of these patterns, for each subject we calculated mean FC values for 6 different groups of cortical connections and compared them using multiple paired sample t-tests **(Figure 3b; see Methods section 7)**. In both hemispheres, FC between core MD regions was significantly stronger than both their connectivity with the penumbra (left *t*(448)= 93.1, right *t*(448)= 79.4), and the internal penumbra connectivity (left *t*(448)= 79.4, right *t*(448)= 66.3). For both core and penumbra MD areas, mean FC with non-MD cortical areas was near zero.

**Figure 3.**
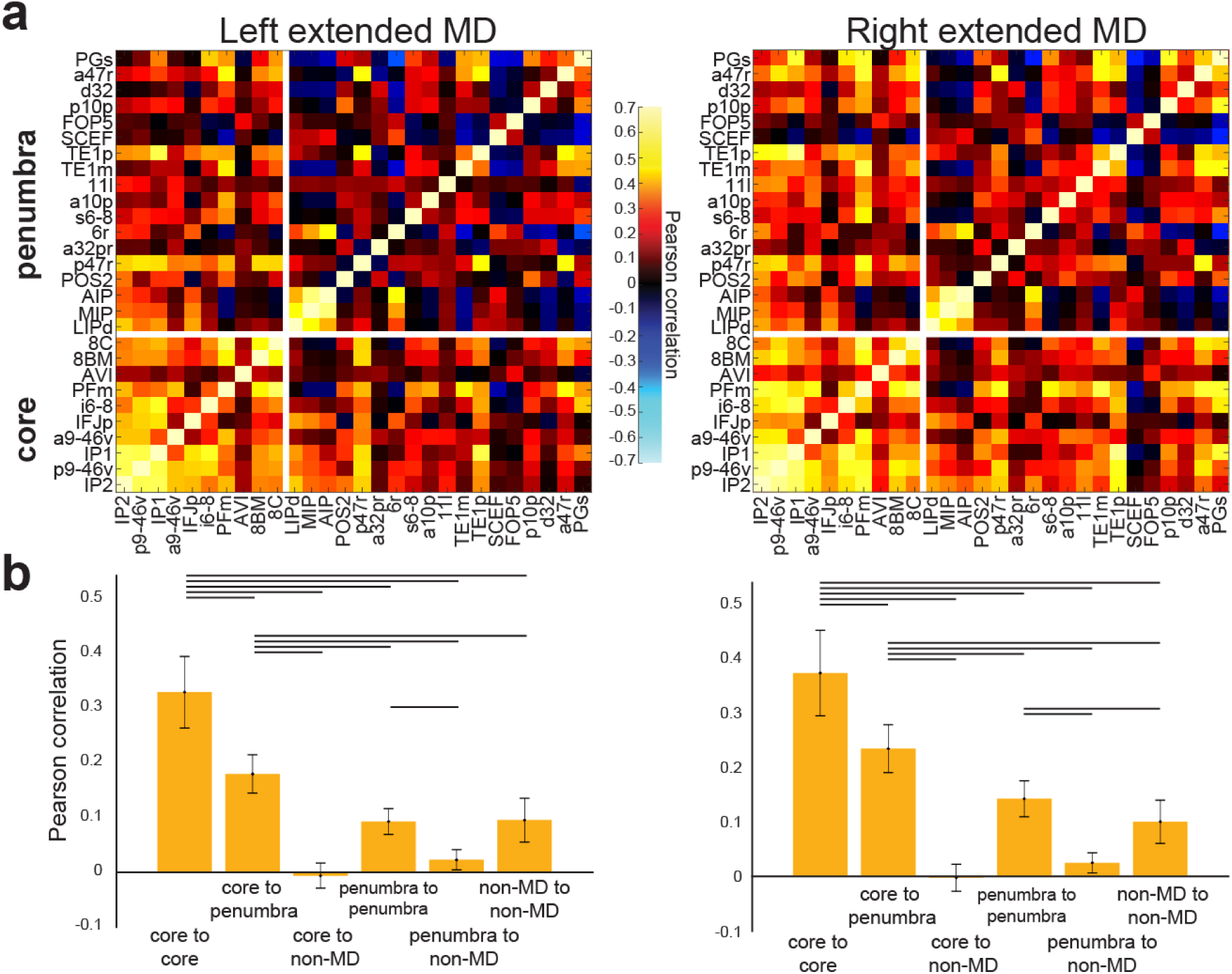
Functional connectivity (FC) of the MD system. **(a)** FC (Pearson correlation) across the MD system. Regions of the extended MD system are separated into core and penumbra, with regions within each set ordered by mean activation (beta) across our 3 functional contrasts. Note the strength of core MD connectivity (lower left box) vs penumbra connectivity (upper right box). **(b)** Statistical comparison (paired sample t-test) between different groups of cortical connections. Lines highlight a statistically significant difference (p<0.05, Bonferroni corrected for 30 comparisons). Data available at http://balsa.wustl.edu/jjL1x

We next investigated the spatial similarity between the MD network defined from our conjunction of 3 task contrasts and canonical fMRI resting state networks. For this purpose, we utilized the recent Cole-Anticevic Brain Network Parcellation (CAB-NP), which analyzed resting state data from 337 HCP subjects and identified network communities across HCP MMP1.0 areas (Ji et al. 2019). A comparison of the extended MD and the CAB-NP (**Figure 4a)** indicates points of both convergence and divergence.

**Figure 4.**
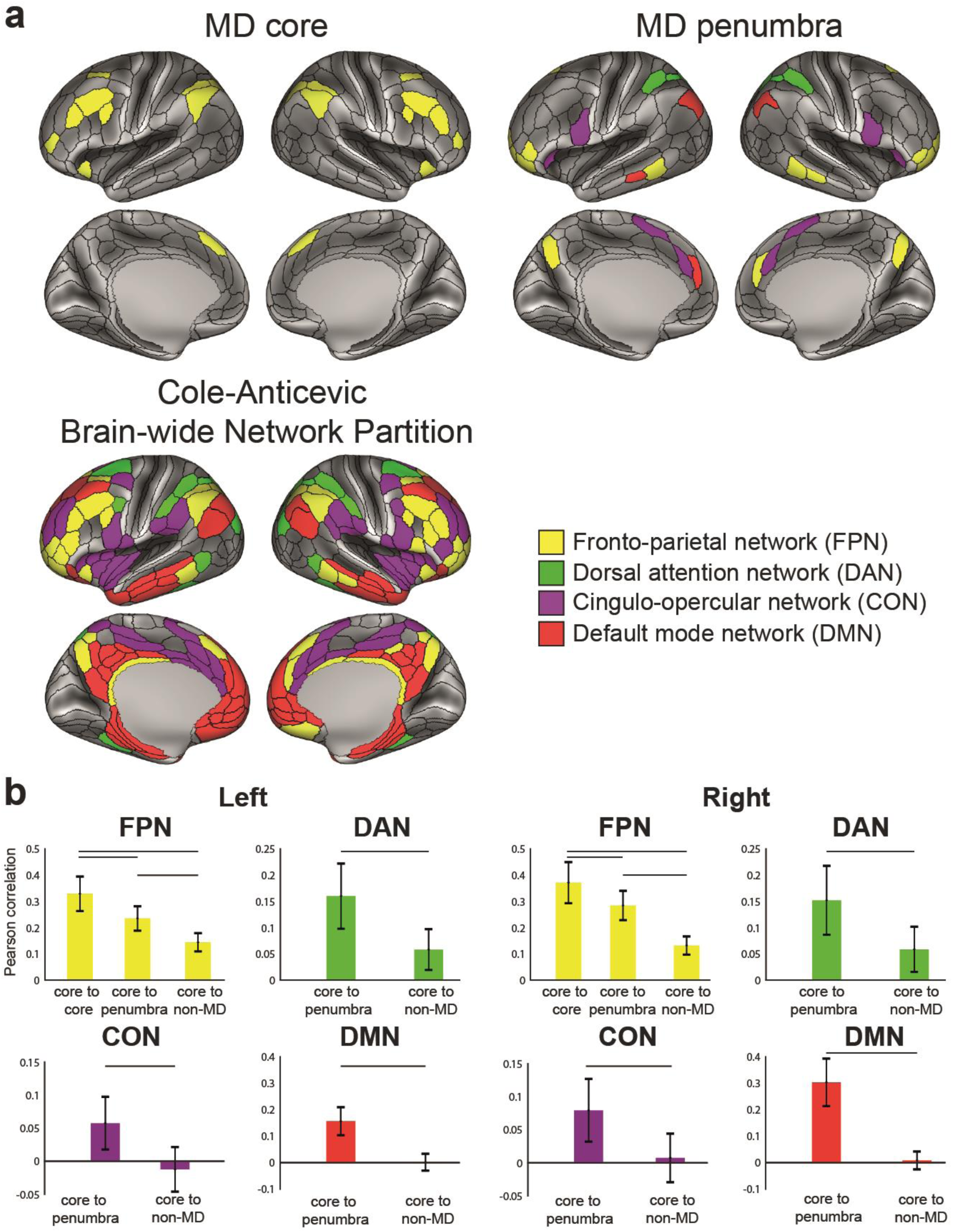
MD system and resting state networks **(a)** Resting state network assignments from the Cole-Anticevic Brain-wide Network Parcellation (CAB-NP; Ji et al., 2019) for the core (top left) and penumbra (top right) MD areas, compared to the whole CAB-NP fronto-parietal network (bottom left). **(b)** Statistical comparison (paired sample t-test) of cortical connection types for each CAB-NP network. Data available at http://balsa.wustl.edu/wNGV6

Most strikingly, all 10 core MD areas are within the fronto-parietal network (FPN), (**Figure 4a**, **top left**). In contrast, penumbra MD areas are scattered among four networks: several in the FPN (yellow, 8 on the left and 10 on the right), 4 in the cingulo-opercular network (CON, purple), 3 in the dorsal attention network (DAN, green) and several in the default mode network (DMN, red; 3 on the left and 1 on the right) **(Figure 4a, top right).** Importantly, examination of the whole CAB-NP FPN network (total 28 areas right, 22 left) shows most but not all areas within the MD core or penumbra (right FPN: 10 core, 10 penumbra, 8 non-MD; left FPN: 10 core, 8 penumbra, 4 non-MD) **(Figure 4a, bottom)**.

To emphasize the central role of core MD, we again compared different connectivity subgroups (**Figure 4b**; paired sample t-tests, p<0.05, Bonferroni corrected). Within the FPN, we found that core MD regions have significantly stronger FC with other FPN regions (core-core vs core-penumbra: left *t*(448)=53.3, right *t*(448)=46.8; and core-core vs core-non-MD FPN regions: left *t*(448)=75.1, right *t*(448)=84.2). Also within the FPN, core-penumbra FC is stronger than core-non-MD FC (left *t*(448)=47.1, right *t*(448)=73.0). We also found higher FC between core MD regions, all within FPN, and penumbra vs non-MD regions within each of DAN, CON and DMN (DAN (left *t*(448)=47.6, right *t*(448)=41.0), CON (left *t*(448)=41.1, right *t*(448)=40.5) and DMN (left *t*(448)=70.1, right *t*(448)=80.4) **(Figure 4b)**.

Many previous studies have separated cognitive control regions into two distinct networks: fronto-parietal (dorso-lateral frontal and intra-parietal sulcus regions) and cingulo-opercular (insular, dorsomedial frontal and anterior lateral frontal regions) (Crittenden et al., 2016; Dosenbach et al., 2008, 2006; Dosenbach et al., 2007; Yeo et al., 2011). We wondered whether our extended MD network would show a similar separation. Multi-dimensional scaling of connectivities between extended MD regions showed core MD regions centrally clustered together (**Figure 5; see Methods section 7)** with no strong trend for a distinct CON among these core regions. Instead, matching their network assignments in CAB-NP, the results suggest a relatively distinct CON cluster including dorsomedial frontal region SCEF and insular region FOP5, These results suggest that the main cingulo-opercular network is distinct from core MD regions, with the two networks in close anatomical proximity.

**Figure 5.**
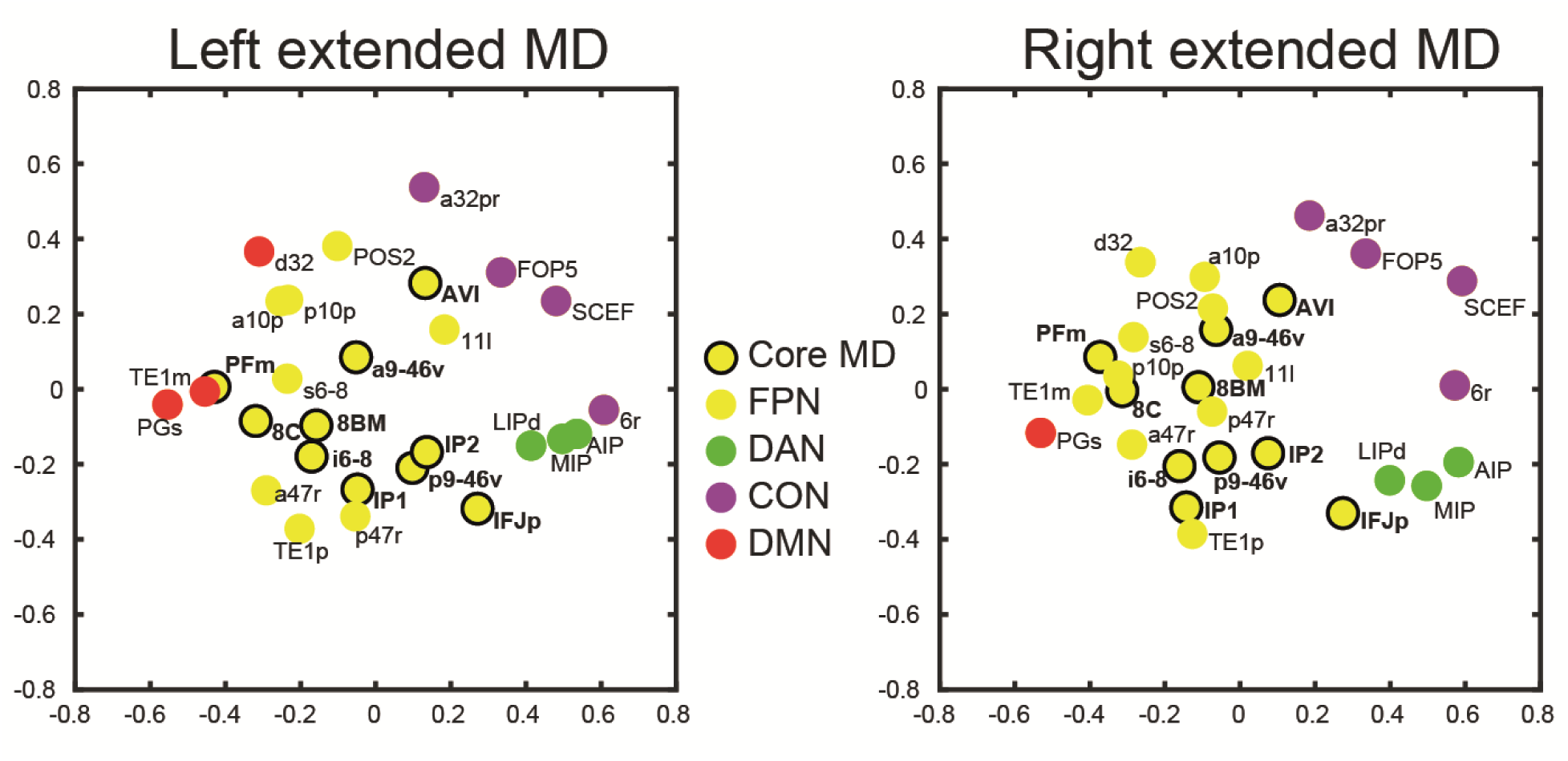
Multi-dimensional scaling plot of the connectivities between extended MD regions. Axis units are arbitrary. Data available at http://balsa.wustl.edu/jjL1x

In summary, while these results show substantial overlap between MD and FPN – especially for the MD core – there are additional organizational aspects revealed by the FC analysis. Connectivity is especially strong between regions within the extended MD system, and strongest between core regions within the canonical FPN. Strong functional connectivity, especially for the core, suggests a suitable architecture for widespread integration of distributed brain states.

Connectivity delineating the MD network can also be revealed by recent work using temporal ICA (tICA), which generates components that are temporally independent (Glasser et al., 2019, 2018; see also Van Essen and Glasser, 2018). By correlating our group average MD map (**Figure 1a)** with the tICA components from (Glasser et al. 2018), we identified at least one rest and one task tICA component having strong spatial similarity to the group average MD map (whole brain absolute Pearson correlation r=0.74 and 0.76 respectively; **Figure S3)**.

### Task profiles across the multiple-demand cortex

By definition, every MD area showed a significant positive result in each of our 3 behavioral contrasts. Across areas, nevertheless, we examined the relative preferences for one contrast over another. To evaluate this quantitatively, **Figure 6a** shows the mean response of each area (averaged across hemispheres) for each contrast. Predominantly, the picture is one of consistency. For nearly all areas, activation was strongest for the Math>Story contrast, and weakest for Relational H>E contrast. Against this general background, however, there was also differentiation between profiles, with varying patterns of peaks and troughs.

**Figure 6.**
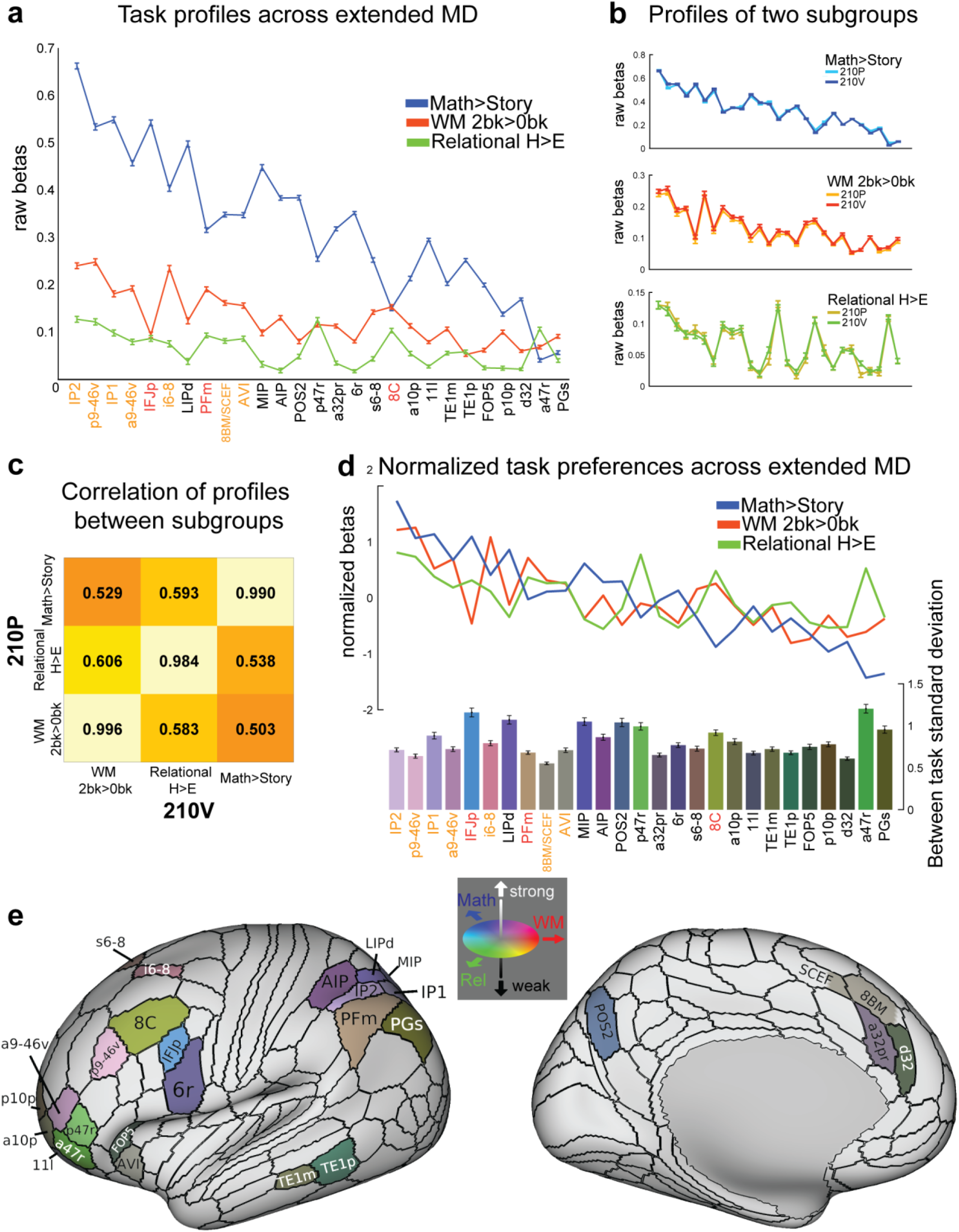
Task profiles across the MD system. **(a)** Raw activation estimates (betas) for each contrast. Areas are sorted from left to right according to the strength of their MD response (average across the 3 contrasts). Error bars represent SEM. Core MD areal labels are colored in orange (survived in all 3 contrasts) and red (survived in 2 out of 3 contrasts). **(b)** Task profiles for two independent groups of subjects (210P and 210V). **(c)** Correlation of task profiles between groups. **(d)** Normalized task profiles across the MD system as line plots. Bar heights represent between-task standard deviation, separately calculated for each subject and averaged over subjects. Bar colors indicate relative preferences between tasks. Color wheel indicates red for working memory (WM), green for relational reasoning (Rel), and blue for math. Intermediate colors show mixed preferences. Brighter and darker colors reflect stronger and weaker MD activation, respectively. **(e)** Cortical projection of the RGB color weighted normalized task profiles. Data available at http://balsa.wustl.edu/4m747

To test the robustness of these patterns, we compared activation profiles in the two independent groups of subjects (210P and 210V). As shown in **Figure 6b**, the activation profile for each contrast is almost identical for the two groups. **Figure 6c** quantifies this by correlating activation profiles **(in Figure 6b)** for the two subject groups. Very high correlations on the diagonal (r > 0.98) highlight how the precise pattern of activation for a given contrast is very stable when averaged over many individuals. Off-diagonal correlations are much lower (r=∼0.5-0.6). A closely similar pattern was seen when extended MD regions were defined in the 210P subgroup, and correlations computed between two halves of the 210V subgroup (diagonals *r*>0.94, off-diagonals *r*=0.25-0.60). Although all tasks engage all MD areas, there remains considerable and highly consistent inter-areal diversity in precise activation patterns.

To illustrate this inter-areal diversity between the three contrasts, we plotted the normalized profile for each contrast **(line plots in Figure 6d).** For each contrast and each subject, we z-scored activations across MD regions, then averaged the z-scores across subjects. For each region, bar heights **(Figure 6d, bottom)** show the standard deviation of these normalized z-scores across tasks, separately calculated for each subject and then averaged over subjects. Bars were also colored to highlight the relative task preferences (see **Figure 6e**, where the same colors are projected onto the cortical surface).

The results reveal a diversity of relative task preferences across the extended MD network. Relative preference for relational reasoning (green) occurs in a cluster of anterior frontal areas inferior to the core region a9-46v, as well as in 8C. Dorsal frontal regions (e.g. i6-8 and s6-8) show relative preference for working memory, whereas dorsal parietal regions (AIP/LIPd/MIP, and POS2) show relative preference for math. Other relative preferences occur across most regions.

Task preferences were also present across hemispheres (**Figure S4**). Most MD regions showed stronger activations in the right hemisphere for both working memory and math contrasts, with more variable results for relational reasoning. Across both hemispheres, however, almost all contrasts were positive, in line with a pattern of largely bilateral MD activation.

Despite relative consistency across the entire extended MD network – with the strongest activation for Math>Story, and weakest for relational reasoning – there is also clear evidence of relative functional specialization, with each area showing modest but consistent relative preference for one contrast over another.

### Multiple-demand regions during weak cognitive demands

A potential limitation of our main analysis is that we might have missed MD regions that are already active even in easy task conditions, and therefore absent in our task contrasts. To investigate this, we examined the group average maps for weak cognitive demands compared against periods during which subjects visually fixated on a cross hair in the middle of the screen. We used the 0bk WM versus fixation contrast (0bk>fix) and easy relational reasoning versus fixation contrast (Relational E>fix) **(Figure 7)**. The Math task did not include any fixation periods and was thus excluded from this analysis.

**Figure 7.**
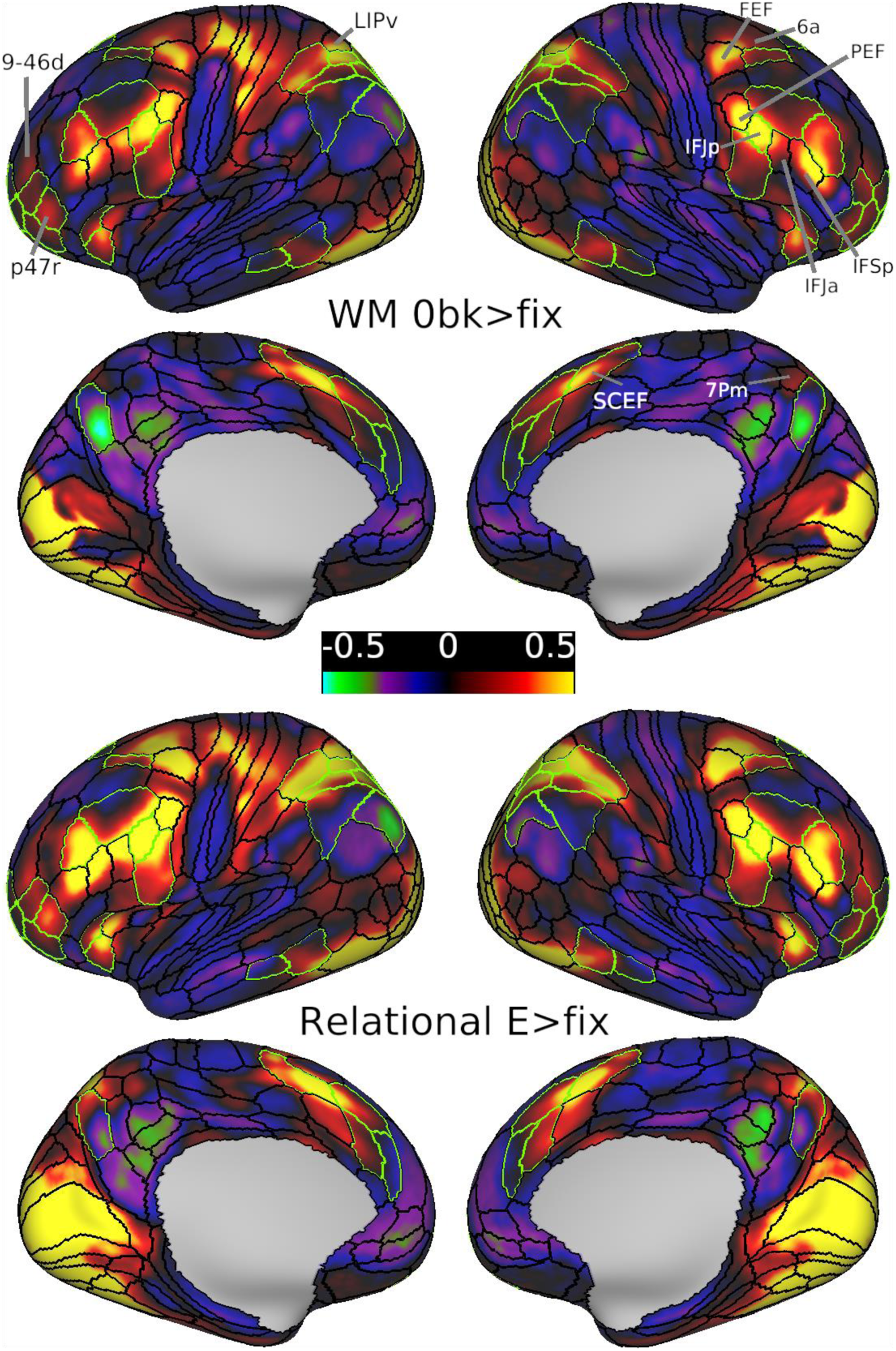
Group average beta maps for the WM 0bk>fix contrast (upper) and Relational E>fix contrast (lower). The borders of extended MD regions are colored in green. Data available at http://balsa.wustl.edu/mDkgN

As expected, the activated regions overlap substantially with the extended MD network, but also include visuo-motor regions as predicted when contrasting task with fixation. In comparison to our previous group MD map **(Figure 1)**, however, there are shifts in the easy vs fixation maps. Dorso-lateral frontal activation shows a posterior shift, with strong activation near the intersection of FEF, 6a and i6-8 areal borders. Premotor frontal activation is strongest around IFJp, spreading towards the premotor eye field (PEF) area dorsally and inferior frontal sulcus regions (IFJa and IFSa) ventrally. Frontal pole activation peaks within penumbra region p47r and also weakly engages area 9-46d in addition to previously identified adjacent MD regions. Dorso-medial frontal activation is strongest within the anterior half of SCEF, spreading anteriorly into 8BM. Lateral parietal activations are strongest around penumbra regions AIP, LIPd and MIP and the adjacent LIPv. Dorso-medial parietal activation overlaps with 7Pm sparing POS2. All previously mentioned regions as well as all core MD regions (except for PFm) were significantly activated in both 0bk>fix and Relational E>fix contrasts (p<0.05; Bonferroni corrected for 180 regions after averaging across hemispheres).

The comparison with fixation-only periods limits the interpretation of activation in the above highlighted regions, as visuo-motor related activation presumably dominates the pattern. For example, activation in FEF and PEF may largely reflect eye movements, especially in the relational task. Tentatively, however, these results suggest that our main task contrasts may miss additional MD regions, extending from those identified in the main analysis, but with strong activation even in the easier version of each task.

### Subcortical and cerebellar components of the multiple-demand system

To identify subcortical and cerebellar components of the MD system we used the same 3 behavioral contrasts used for cortical areas. FreeSurfer’s standard segmentation of 19 subcortical/cerebellar structures (left and right caudate, putamen, globus pallidus, thalamus, cerebellum, hippocampus, amygdala, ventral diencephalon, nucleus accumbens; plus whole brainstem) was carried out separately for every subject (see **Methods section 4**), thus avoiding mixing signals from nearby structures or white matter. For each structure, we first identified significantly activated voxels for each contrast separately (one sample t-test, FDR corrected for each structure separately, p<0.05, Bonferroni corrected for 19 structures) and then identified the conjunction of significant voxels across the three contrasts. We analyzed the P210 and V210 groups separately. This revealed activation regions bilaterally mainly in the caudate nucleus and cerebellum. Caudate activation was in a circumscribed region in the head, which was modestly replicable between 210V and 210P groups (*r*=0.37, Dice=0.60 across all caudate voxels) **(Figure 8a, left panel)**. Cerebellar activations, mapped to a cerebellar surface model (Diedrichsen and Zotow 2015) and displayed on a cerebellar flatmap, included separate medial and lateral portions of crus I and II (on dorsal and ventral lateral surface). The pattern was largely symmetrical across hemispheres and was strongly replicable across both groups (*r*=0.88, Dice=0.88) **(Figure 8a, right panel)**.

**Figure 8.**
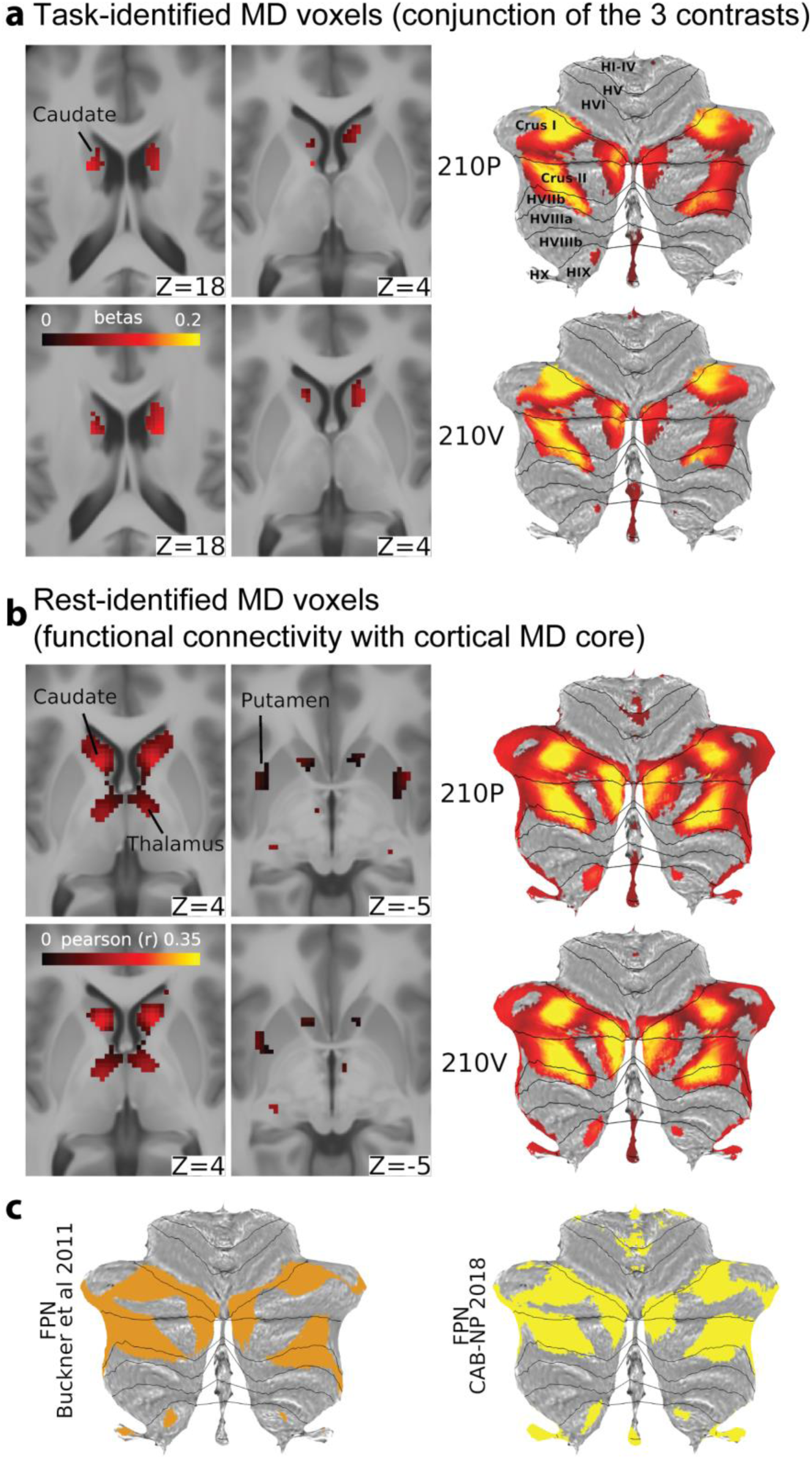
Subcortical and cerebellar MD components. **(a)** Left: Conjunction of significant voxels across the three tasks for the 210P (top) and 210V (bottom). Right: Cerebellar activity is displayed on a flat cerebellum with lines representing anatomical borders (Diedrichsen and Zotow 2015). Data available at http://balsa.wustl.edu/Z4NXp **(b)** Left: Subcortical voxels with significant connections to the cortical core MD areas. Right: Cerebellar MD connectivity displayed on a flat map. Data available at http://balsa.wustl.edu/VjwZg **(c)** FPN from Buckner et al. (2011) (left) and Ji et al. (2019) (right). Data available at http://balsa.wustl.edu/3g7wv

The analysis showed no significant regions in the thalamus, putamen or globus pallidus **(Figure 8a)**. Interestingly, larger bilateral portions of the thalamus (anterior dorso-medial), putamen (dorso-anterior/mid portion) and globus pallidus (dorso-anterior portion) were significantly activated in only two contrasts (working memory and math) and were deactivated in the relational reasoning contrast **(Figure S5)**.

In a separate analysis using resting state data, we aimed to identify the subcortical and cerebellar voxels showing significant functional connectivity with the cortical core MD areas. For this analysis we used the group average dense FC matrix for each group **(see Methods section 7). Figure 8b** shows the statistically significant subcortical/cerebellar voxels (FDR corrected, p<0.05, Bonferroni corrected for 19 structures). The patterns were highly replicable (caudate *r*=0.84, Dice=0.94; cerebellum *r*=0.97, Dice=0.93) and follow closely the task-identified regions in the caudate nucleus and cerebellum bilaterally. In addition, FC analysis identified significant voxels in bilateral portions of the thalamus (anterior dorso-medial) and putamen (dorso-anterior/mid portion), similar to the regions activated in the working memory and math contrasts **(Figure S5)**. We also note that a similar overlapping thalamic region is activated in the Relational E>fix contrast **(Figure S5).**

We also compared the MD cerebellar regions with the fronto-parietal network (FPN) identified by resting state data from two studies: Buckner et al. 2011 (7 networks parcellation results from 1000 subjects) and CAB-NP (Ji et al., 2019; results from 339 HCP subjects). **Figure 8c** illustrates the strong similarity between the FPNs from both studies and the cerebellar MD hotspots in crus I and II (Dice=0.62-0.70).

Next we measured the similarity between the task and rest identified subcortical and cerebellar MD regions (after conjunction of 210P and 210V maps). With the exception of the left caudate, task and rest fMRI data showed modest overlap (left caudate *r*=0.01, Dice=0.07; right caudate *r*=0.18, Dice=0.26; left cerebellum *r*=0.65, Dice=0.68; right cerebellum *r*=0.60, Dice=0.62). Thus, together, task and rest fMRI data converge on identifying subcortical, especially caudate, and cerebellar regions related to the cortical MD core.

## Discussion

Thousands of brain imaging studies have identified regions of frontal and parietal activation crossing multiple cognitive demands. In this study, we used HCP’s high quality multimodal MRI data and improved brain registration methods to demonstrate that diverse cognitive tasks from different sensory modalities engage widely distributed but tightly delineated foci of multiple-demand (MD) activation **(Figure 1a)**. The network of twenty-seven extended MD areas is organized into nine larger patches **(Figure 1a, b)**: three distributed in an anterior-posterior chain running along the lateral frontal surface, a fourth in and above the anterior insula, a fifth on the most dorsal part of the lateral frontal surface, a sixth on the dorsomedial frontal surface, a seventh within and surrounding the intraparietal sulcus, an eighth in the dorsomedial parietal cortex, and a ninth in posterior temporal cortex. Within these larger patches, we identified a set of core areas, characterized by their strong activation and FC-based interconnectivity, surrounded by a penumbra having relatively weaker activations and interconnectivity. We also identified localized MD regions in the caudate nucleus and cerebellum that share strong connectivity with the cortical core MD. These data provide strong evidence for the existence of highly specific MD regions in the human brain. The improved anatomical precision offered by the HCP methods revealed novel findings regarding the anatomical and functional organization of the MD network, as well as the functional connectivity of its components.

Why should the brain contain this precise network of MD regions, co-activated during many cognitive activities? Within the extended MD system, we propose that the core regions, most strongly active and interconnected lie at the heart of information integration and exchange mediating cognitive operations. Surrounding penumbra regions, with their connectivity into multiple cortical networks, feed diverse information into and out of the core. Across the entire MD system, co-activation reflects rapid information integration and exchange, while modest functional preferences reflect differential connectivity and information access. Together, these properties allow MD regions, with associated subcortical regions, to build integrated cognitive structures suited to current behavioral requirements. These proposals are developed and extended in the following sections.

### Broad anatomical distribution and relative functional preferences

Similar activation patterns crossing many cognitive domains, roughly corresponding to our current MD findings, has been documented in a large body of previous work. At the same time, there have been many suggestions of functional differentiation between MD-like regions, albeit with little consensus emerging across studies (Dosenbach et al. 2007; Champod and Petrides 2010; Hampshire et al. 2012; Yeo et al. 2015; Lorenz et al. 2018). Our fine-grained anatomical findings illustrate the challenges in interpreting studies that are based on traditional neuroimaging analyses. For example, when coarsely-analyzed data suggest functional dissociation between MD-like regions, the dissociation might concern penumbra or core MD regions, or even nearby non-MD regions that are more task specific. (See **Figure S2** for task specific activations for each of the 3 contrasts extending beyond MD parcels; also see Coalson et al. 2018 for a quantification of the uncertainties involved in mapping between volumetric activations and surface activations.) The finer-grained anatomy of the current study helps clarify issues of functional differentiation within the MD system. On one side is strong evidence for a network of co-activated MD regions, broadly distributed across the cortex. On the other is strong evidence for relative functional differentiation, often somewhat corresponding to previous proposals in the literature. Below we summarize concrete functional questions that are clarified by the present data.

Much prior work (see **Figure 1b**) has suggested MD-like activation in the posterior dorsal prefrontal cortex, in a region close to the FEF. Though a reasonable interpretation might be increased eye movements in more demanding conditions, we show that the main focus of MD activation is localized anterior and dorsal to the FEF, including regions i6-8 (core) and s6-8. These results strongly suggest that MD activation is distinct from activations driven simply by eye movements in complex tasks. Our results match an early demonstration of working memory activation immediately anterior to FEF (Courtney et al. 1998); in our data, the strong MD response of i6-8 and s6-8 is supplemented by relative preference for the working memory contrast (see **Figure 5d**).

Near the frontal pole, we localized MD activation in one core region (a9-46v) and 5 surrounding penumbra regions. There has been much debate concerning an anterior-posterior gradient of activation on the lateral frontal surface. On the one hand, many tasks produce activation near to the frontal pole, suggesting an MD-like pattern (Ramnani and Owen 2004). On the other hand, many studies suggest selective activation in this region, for example associated with abstract reasoning (Bunge 2004; Christoff et al. 2009) or hierarchically-organized cognitive control (Badre 2008; Badre and Nee 2018). Our results indicate that a9-46v is almost as strongly co-activated as more posterior core regions, arguing against a simple gradient of activation. Its adjacent penumbra regions (a47r, p47r) also show clear MD activation but with relative functional preference for the abstract relational reasoning task, matching previous reports of reasoning activation in this region.

The combined 8BM/SCEF MD area on the medial frontal surface showed the least functional preference **(Figure 5d)**. Our findings show MD activation rising to and peaking at the border between 8BM and SCEF, with similar patterns also visible in other task contrasts and fine-grained analysis of functional connectivity **(Figure S2).** In our group-average map, hints of peak task activation near areal borders can also be seen at the borders of 8C/IFJp and POS2/7Pm **(Figure 1a)**. Though detailed analysis of these functional transitions is beyond our scope here, it is possible that here too MD activation peaks near areal borders. Borders between these areas were defined using robust multiple overlapping functional, architectural and/or topological criteria (Glasser, Coalson, et al. 2016). Thus, we speculate that our data may reflect close interaction between areas sharing a common border, reflecting the general principle of spatial proximity between brain regions that are in close communication.

Previously, many studies have reported a band of occipito-temporal activation accompanying activation of fronto-parietal MD regions **(see Figure 1b)**. As most tasks used in these studies were visual, a plausible interpretation might be top-down input into higher visual areas. In our data we identified two penumbra regions, TE1m and TE1p, in posterior temporal cortex. Since these regions were activated by the auditory as well as the visual contrasts, the interpretation of top-down input into higher visual areas is less plausible. The location of these regions midway between higher visual areas, auditory areas and language and semantic areas (Pobric et al. 2007; Visser et al. 2010; Fedorenko et al. 2011) suggests a genuine MD region, situated to integrate higher visual, auditory and semantic/language processing. Similar to previous findings in Broca’s area (see Fedorenko et al., 2012), these data highlight an MD area with close proximity to language regions.

Previous studies employing math tasks identify an MD-like pattern that is commonly interpreted as a domain-specific “math network” (Amalric and Dehaene 2017). Our results show that the math contrast engages all extended MD regions, but with relative preferences among dorsal parietal areas (AIP, LIPd, MIP; and POS2 on the medial surface) and dorsal frontal region IFJp. We note that in our data, math preferences are potentially confounded with auditory preferences (Michalka et al. 2015).

Our selected task contrasts might have led us to miss MD regions that were already active in the easier tasks. Indeed, comparison of easy tasks with fixation suggested extension of MD activation into adjacent regions, including FEF, PEF, 9-46d, 7Pm and LIPv. Evidence that even easy tasks produce strong activation in posterior regions of the lateral frontal cortex fits a number of previous reports (Badre 2008; Crittenden and Duncan 2014; Shashidhara et al. 2019). At present, the limited number of suitable contrasts in the HCP data and the difficulty of interpreting contrasts with fixation preclude strong conclusions on these additional putative MD regions. For example, while activation of FEF and PEF might simply reflect eye movements, this interpretation could be incomplete given that one easy task (0bk) used only stimuli placed in the center of the visual field. Future studies utilizing HCP methods and examining a broader range of task contrasts should provide clearer answers.

In line with much current thinking, relative functional specializations might suggest that different MD regions are specialized for different cognitive operations. Though this interpretation is reasonable, it leaves open the question of why these regions are active in such a diversity of tasks, how they communicate and coordinate their activities, why their representations show such flexibility, and why they have such consistently strong functional connectivity. Instead of strong functional specialization, we suggest that distributed MD regions serve to combine and relate the multiple components of cognitive operations. While data from macaques show that putative MD regions share many anatomical connections (Selemon and Goldman-Rakic 1988; Mitchell et al. 2016), each also has its own unique fingerprint of connections to and from other brain regions (Petrides and Pandya 1999; Markov et al. 2014). Thus the wide distribution and diverse connections of MD regions likely provides the necessary anatomical skeleton for access to different kinds of information from different brain systems. Different tasks, emphasizing different kinds of information, then lead to quantitatively different patterns of activation across the MD system. At the same time, rich interconnections between MD regions allow information to be rapidly exchanged and integrated.

To extend the present results, a wider range of task contrasts would be valuable. Though the 3 contrasts used here are already quite diverse, a wider range of contrasts could establish boundary conditions on MD recruitment, and add more detailed understanding of relative functional preferences. One open question concerns strong manipulations of cognitive demand that produce little MD activation. Most conspicuously, some studies (e.g. Han and Marois 2014; Wen et al. 2018) – though certainly not all (e.g. Jiang and Kanwisher 2003; Crittenden and Duncan 2014) – suggest little MD activation for demanding visual discriminations limited by the quality of sensory data. Though we would contend that any task requires integration of its components, we might speculate that integration demands do not limit performance in simple sensory tasks. Such exceptions to the MD pattern remain an important topic for future work.

### MD cortex and resting state networks

In this study we identified the extended MD system using a conjunction of three task contrasts. Using MD regions identified from task data, we proceeded to demonstrate strong within-network functional connectivity at rest. As expected, our analysis of resting state data shows much convergence with canonical functional networks derived from the same data (Ji et al. 2019), but we also found additional fine-grained structure. MD core regions constitute a subset of areas within the canonical FPN that are distinguished by especially strong mutual connectivity. This strong connectivity includes widely separated areas. In contrast to the MD core, penumbra regions are distributed across four canonical networks. Again, compared to other regions within those networks, they are distinguished by especially strong connectivity with the MD core. These results support the picture of MD regions as a strong communication skeleton, with penumbra regions in particular drawing together information from several distinct large-scale networks.

In some prior work (Dosenbach et al. 2006, 2007, 2008), insula, dorsomedial frontal and anterior lateral frontal regions have been combined into a CON network, separate from other control regions forming the FPN. In line with CAB-NP, our precise delineation suggest a slightly different picture, with specific regions of anterior insula (AVI), dorsomedial frontal (8BM) and anterior lateral frontal cortex (a9-46v) included in the MD core, and closely adjacent regions included in a separate CON.

Our conclusions are reminiscent of extensive recent work using network science approaches (e.g., graph theory) to identify putative cortical communication hubs (Sporns 2014; Petersen and Sporns 2015; Bassett and Sporns 2017; Bertolero et al. 2018). In this graph theoretic approach, hubs are defined by broad connectivity and/or spatial proximity with multiple cortical networks. Typically they include a set of regions resembling the current MD system, but also others including the temporo-parietal junction, extensive regions of the mid- and posterior cingulate and more (Power et al. 2013; Gordon et al. 2018). These connectional findings are broadly consistent with our proposal that MD regions act as an integrative skeleton for cognitive activity, but they leave open the question of precise relations between the MD pattern, defined with converging task contrasts, and the definition of hubs based solely on functional connectivity. Because hubs are defined by connectivity with multiple cortical networks, their identification depends on the granularity with which these networks are separated and by other factors, including the threshold used to define network ‘edges’ and by potential methodological biases that are commonly overlooked, such as regional differences in receive coil sensitivity that may impact FC values. Such limitations do not apply to definition of MD regions based on converging task contrasts. Further work may help to contrast the functional role of MD regions relative to hubs defined by connectivity but not showing robust activation across multiple diverse tasks.

### Subcortical and cerebellar MD regions

We found MD activation and strong functional connectivity with the cortical MD core in the head of the caudate nucleus. In nonhuman primates, the anterior portion of the caudate receives projections from all prefrontal regions (Averbeck et al. 2014). Tracer studies have established that the dorso-lateral prefrontal, dorso-medial prefrontal and parietal cortices, in addition to strong mutual interconnections, also share converging projections to the caudate, mainly targeting its head (Kemp and Powell 1970; Alexander et al. 1986; Yeterian and Pandya 1991; Middleton and Strick 2000; Haber 2003; Hampson et al. 2006; Choi et al. 2016). Within the striatum, overlap in the projection zones of nearby cortical areas may in part be mediated by interdigitating dendrites and axons that cross functional boundaries (Haber 2003; Averbeck et al. 2014). These anatomical findings are consistent with the identified MD activations in the head of the caudate and strongly support its putative role in information integration.

We also identified distributed MD regions in the cerebellum. Tracer studies identify polysynaptic connections between the prefrontal cortex and the lateral portions of crus I and II as well as vermal lobules VII and IX (Bostan et al. 2013), largely overlapping with our MD cerebellar regions. In addition, previous studies have implicated similar cerebellar regions in several aspects of complex cognitive activity (King et al. 2019) as well as encoding task-relevant information (Balsters et al. 2013). Importantly, MD cerebellar regions do not overlap with motor-related regions (Diedrichsen and Zotow 2015). Not surprisingly, there is strong overlap between the cerebellar regions identified here using converging task contrasts and strong connectivity with the MD cortical core, and the FPN-related cerebellar network defined in previous studies (Buckner et al. 2011; Ji et al. 2019). Importantly, the cerebellar MD regions were identified by connectivity with the more spatially restricted cortical MD core in comparison with the cortical FPN, further suggesting a central role for the cortical MD core.

Based on resting state connectivity, we also identified putative MD regions in the anterior portion of the thalamus. The connectivity-identified thalamic regions are in line with numerous studies reporting strong anatomical and functional connectivity between thalamic nuclei (especially medio-dorsal portions) and fronto-parietal cortex (Haber 2003; Halassa and Kastner 2017). A similar thalamic region was also identified by the conjunction of working memory and math contrasts; for relational reasoning, however, this thalamic region was already active in the contrast of easy task vs rest, with no further increase in the harder task version.

Further work at higher field MRI strength (e.g., 7T) may help clarify the role of these and other subcortical regions associated with the cortical MD system. Meanwhile, in agreement with known anatomy, our data suggest extensive cortical-subcortical interaction in control of complex cognitive activity.

### A precisely-localized neural system supporting complex cognition

For continued progress in understanding brain functional organization, a basic step is delineation of an accepted set of component regions. In the case of MD activation, progress has been slow because we lack such a precise definition, leading to many thousands of studies showing similar activation patterns, but little agreement over questions such as functional similarity/differentiation. Based on the HCP multi-modal parcellation, our work defines a precise network of core MD regions and their surrounding penumbra, and establishes a pattern of widespread co-recruitment, relative functional differentiation, and strong functional connectivity.

These properties support a central role for the MD system in supporting complex cognition. The richness of even a simple cognitive event, and the precise relations that must be established between its different components, call for a widely-connected system, able to access any kind of cognitive content. Owing to their differential anatomical and functional connections, different MD regions may be preferentially recruited as different cognitive contents are accessed. However, strong interconnection between MD regions likely allows different information to become quickly integrated and exchanged, leading to a dominant pattern of co-activation. Extensive MD connections with other regions also suggest a broad role in coordinating brain activity in service of the task at hand. This proposal conforms with the finding that the MD system, among different brain networks, is the most striking in changing its global brain connectivity during different task states (Cole et al. 2013). For future studies, precisely specified MD regions provide the groundwork for detailed functional analyses, cross-reference between studies, and identification of cross-species homologs. This holds promise for a new and more productive phase in study of this core brain network.

## Acknowledgments

Data were provided by the Human Connectome Project, WU-Minn Consortium (Principal Investigators: David Van Essen and Kamil Ugurbil; 1U54MH091657) funded by the 16 NIH Institutes and Centers that support the NIH Blueprint for Neuroscience Research; and by the McDonnell Center for Systems Neuroscience at Washington University. We thank Michael Cole and Alan Anticevic for sharing their network parcellation data prior to publication. We thank Daniel Mitchell for sharing the MD volumetric map.

## Author Contributions (CRediT taxonomy)

Conceptualization, J.D., M.A.; Methodology, M.A, J.D., M.F.G, D.C.V.E; Formal Analysis, M.A., M.F.G; Writing – original draft, M.A, J.D.; Writing – Review & Editing M.A, J.D., M.F.G, D.C.V.E

