## Supplementary material for "A Domain-general Cognitive Core defined in Multimodally Parcellated Human Cortex": Figure S1, Figure S2, Figure S3, Figure S4

### Supplementary Figures

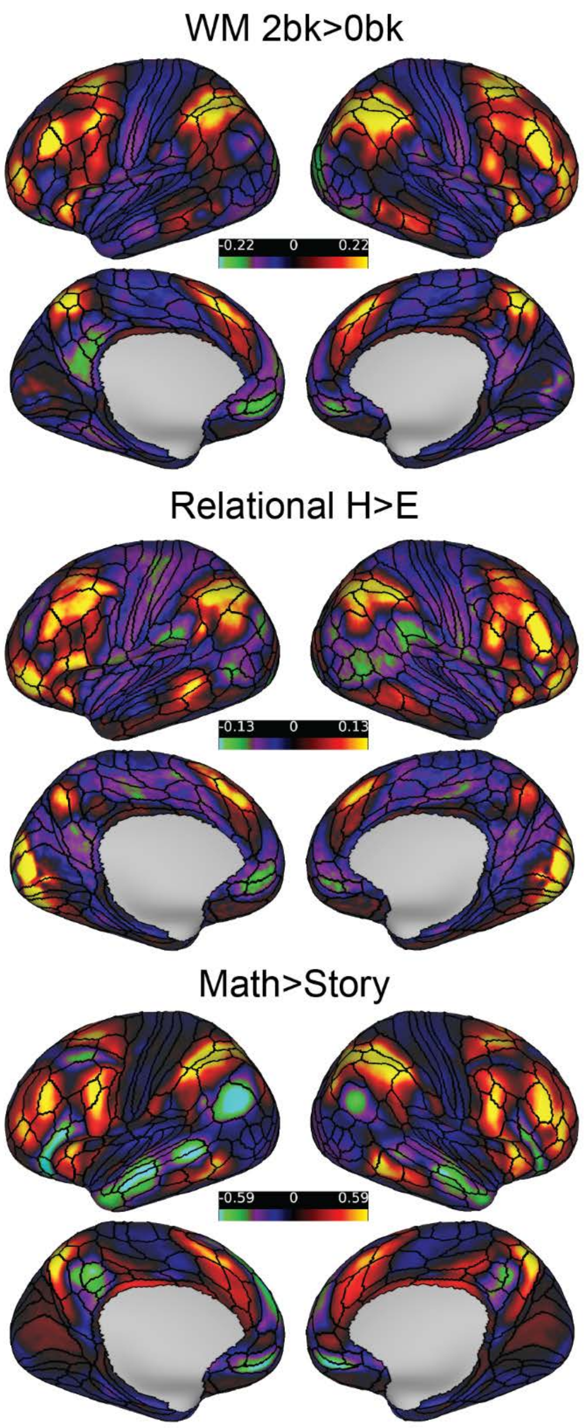


**Figure S1. Contrast maps for each task**. Data available at https://balsa.wustl.edu/zp9XZ

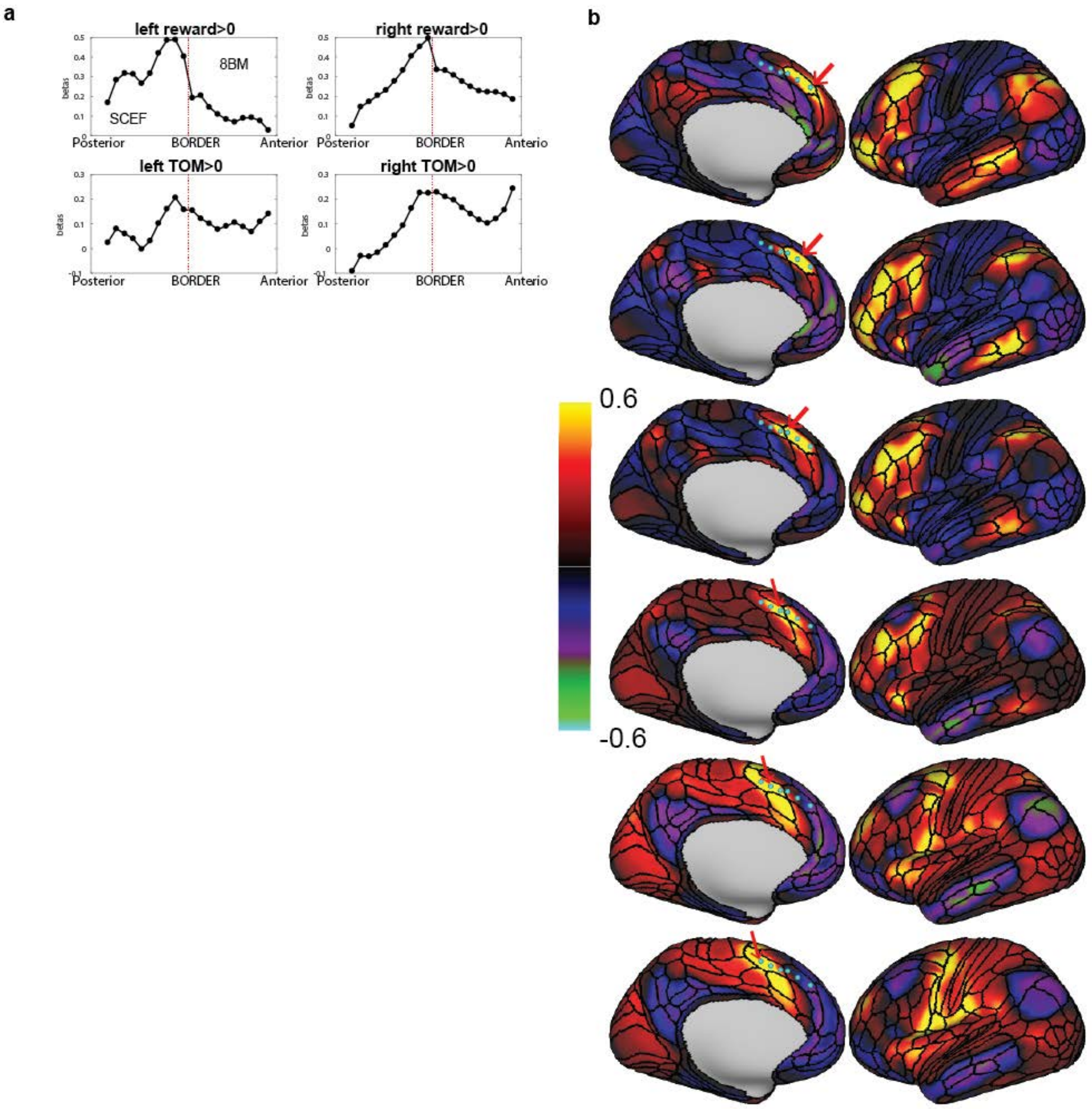


**Figure S2. 8BM/SCEF border. (a)** Group average responses for two HCP contrasts across the 8BM/SCEF border, Reward>fix and Theory of Mind (TOM)>fix, showing a similar pattern of build up within SCEF reaching a peak near the 8BM/SCEF border. **(b)** Functional connectivity maps for seeds (210V map, left hemisphere) along an antero-posterior gradient for the left 8BM/SCEF areas. Arrows mark the seed related to each column's maps. Note how the seed in row 4 is in SCEF near the 8BM/SCEF border and still shows an MD like connectivity pattern, especially the strong connectivity to i6-8. More posterior seeds in SCEF show a markedly different pattern with strong connectivity to FEF. Color scale is Pearson correlation (r). Data available at https://balsa.wustl.edu/X5q36

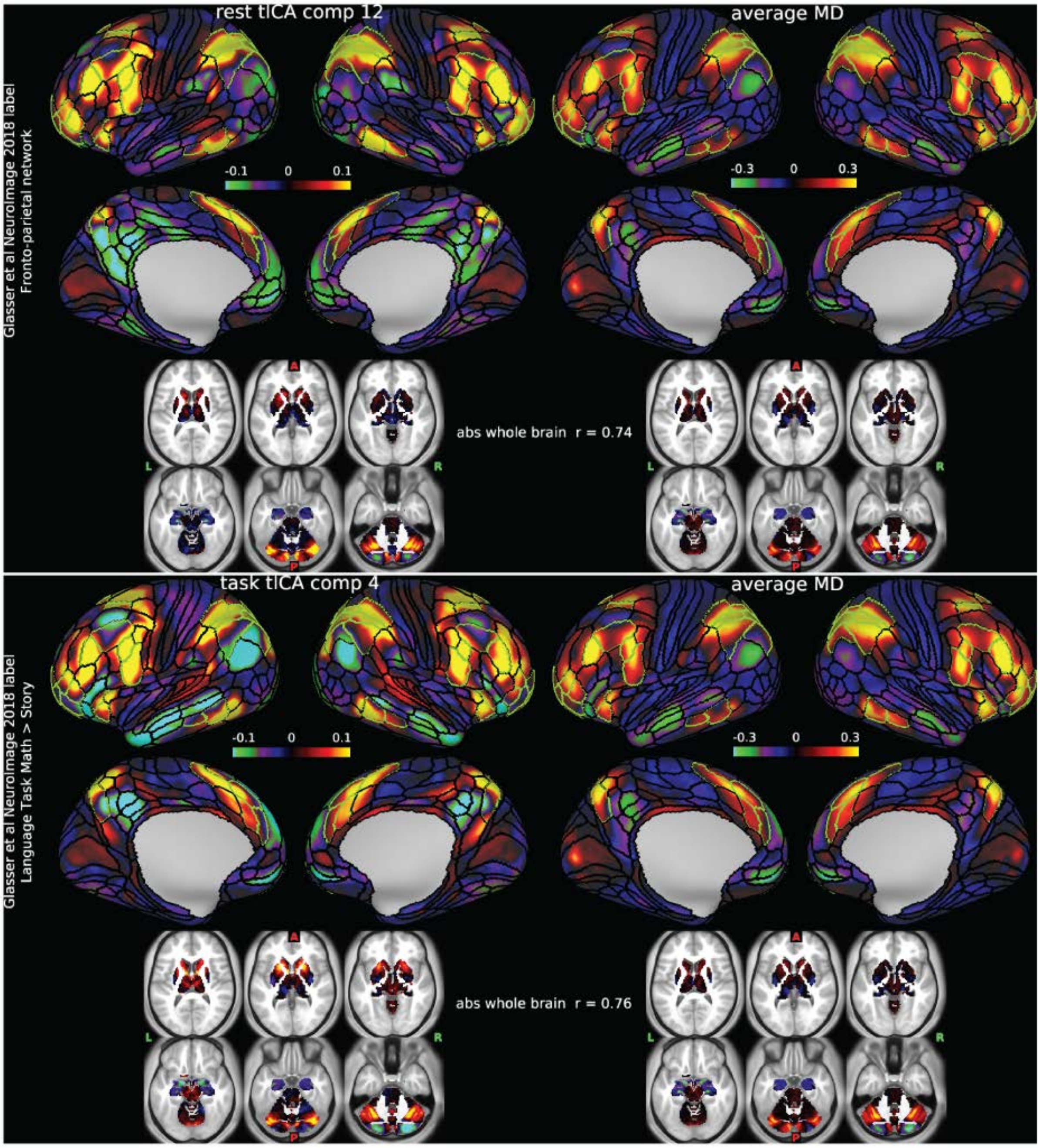


**Figure S3. MD and temporal ICA.** Most correlated temporal ICA components (from Glasser et al., 2018) with MD average map. Top: rest tICA component 12. Bottom: task tICA component 4. The borders of extended MD regions are colored in green. Data available at https://balsa.wustl.edu/87P3x and https://balsa.wustl.edu/Klv5g

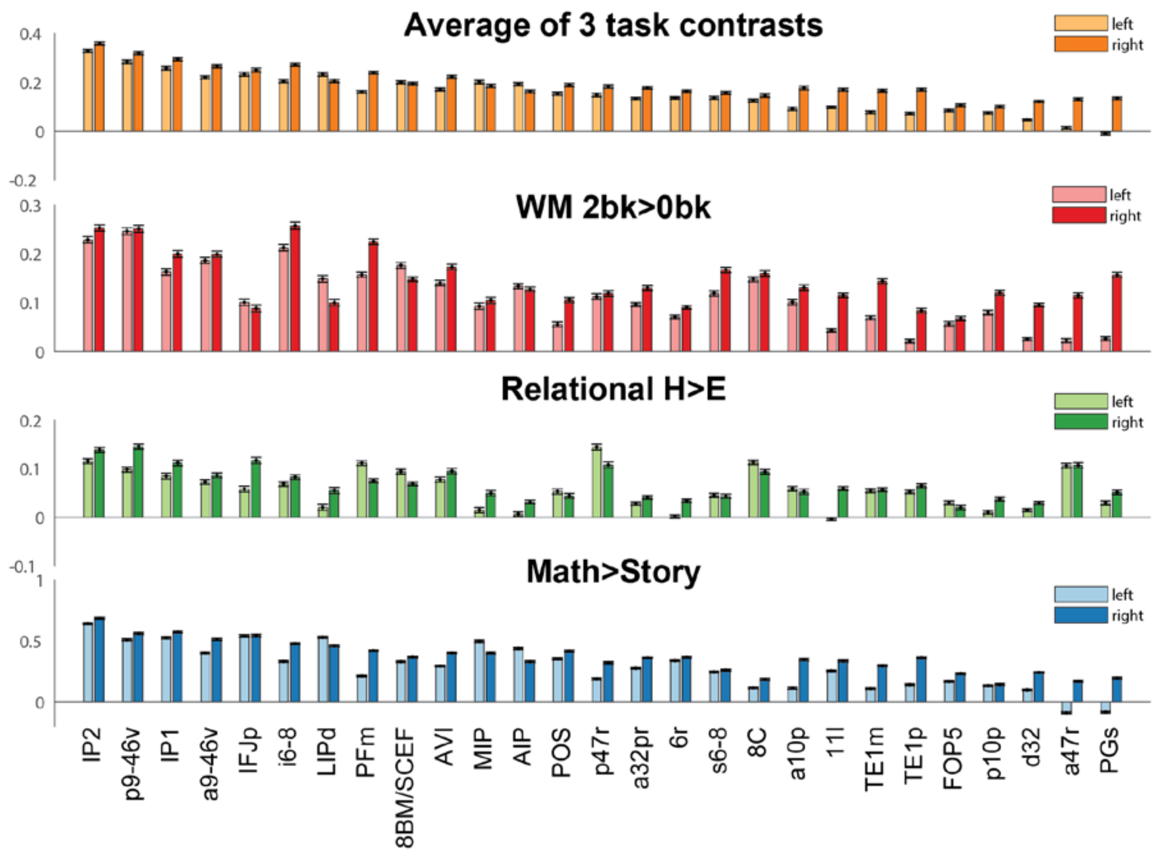


**Figure S4. Extended MD for each hemisphere.** Group average responses for the MD areas of both hemispheres. First row: average of the 3 HCP contrasts. Second row: Working memory. Third row: Relational reasoning. Fourth row: Math>story. Error bars are SEMs.

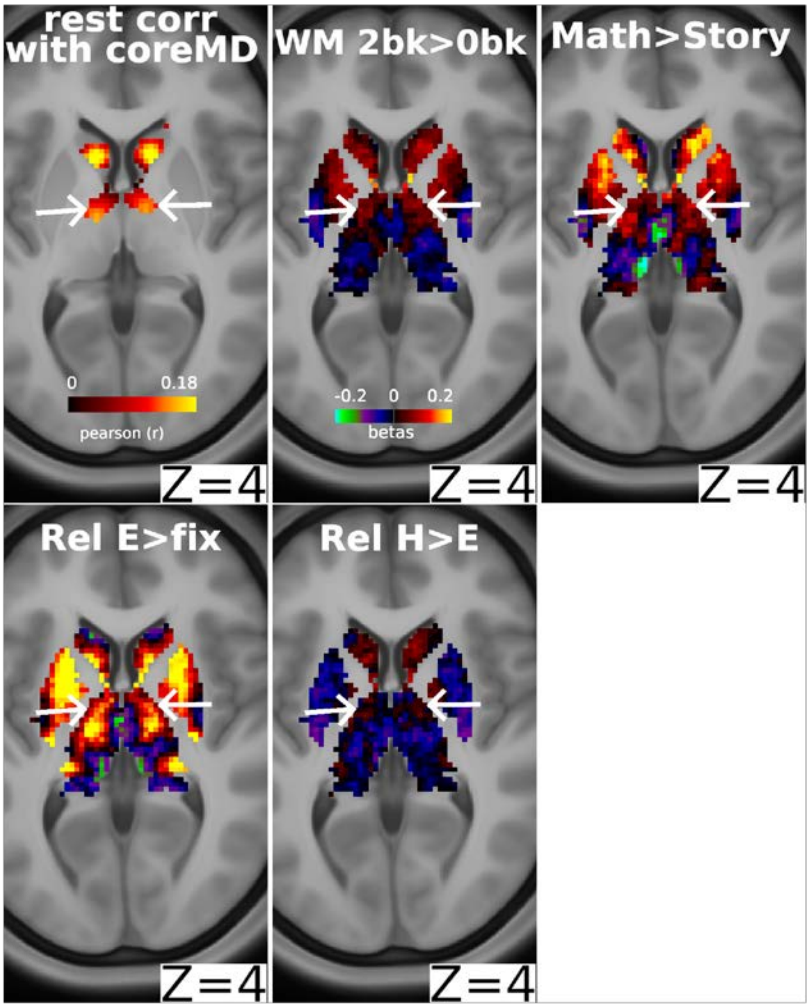


**Figure S5. Subcortical MD.** *Top left*: Subcortical voxels with significant connections to the cortical core MD areas**.** *All other panels*: Group average activity for each task contrast. Arrows highlights that the thalamic hotspot in the top row panels is also activated in the Relational E>fix contrast (*bottom left)*. Data available at https://balsa.wustl.edu/Nw1MK
